# Think zinc: Role of zinc poisoning in the intraphagosomal killing of bacteria by the amoeba *Dictyostelium*

**DOI:** 10.1101/356949

**Authors:** Caroline Barisch, Vera Kalinina, Louise H. Lefrançois, Joddy Appiah, Thierry Soldati

## Abstract

Professional phagocytes have developed an extensive repertoire of autonomous immunity strategies to ensure killing of bacteria. Besides phagosome acidification and the generation of reactive oxygen species, deprivation of nutrients and the lumenal accumulation of toxic metals are essential to kill ingested bacteria or inhibit growth of intracellular pathogens. We use the soil amoeba *Dictyostelium discoideum*, a professional phagocyte that digests bacteria for nutritional purposes, to decipher the role of zinc poisoning during phagocytosis of non-pathogenic bacteria and visualize the temporal and spatial dynamics of compartmentalized, free zinc using fluorescent probes. Immediately after particle uptake, zinc is delivered to phagosomes by fusion with “zincosomes” of endosomal origin, but also by the action of one or more zinc transporters. We localize the four *Dictyostelium* ZnT transporters to endosomes, the contractile vacuole and the Golgi apparatus, and study the impact of *znt* knockouts on zinc homeostasis. Finally, we show that zinc is delivered into the lumen of *Mycobacterium smegmatis*-containing vacuoles, and that *Escherichia coli* deficient in the zinc efflux P_1B_-type ATPase ZntA is killed faster than wild type bacteria.

**Summary statement:** Metal poisoning is one of the bactericidal strategies of macrophages. Here, we describe the dynamics of free Zn and the role of Zn transporters during phagocytosis in *Dictyostelium*.

## Introduction

Transition metals such as iron (Fe), zinc (Zn), manganese (Mn), cobalt (Co) and copper (Cu) are essential for the survival of all living organisms [reviewed in (Weiss and Carver, 2018)]. These metals are incorporated into active sites of metalloenzymes, and organise secondary structures such as Zn finger domains of many proteins including transcription factors. They are therefore implicated in a wide range of crucial biological processes. However, excess of these metals is toxic for living organisms, partially because they compete for metal-binding sites in enzymes. In this context, recent studies have revealed transition metals as a “double edge sword” during infection of phagocytic cells. Upon phagocytosis, the innate immune phagocyte restricts intravacuolar bacterial growth either by depleting essential metal ions (e.g. Fe^2+^ and Mn^2+^) or by accumulating others, such as Cu^2+^ and Zn^2+^, to intoxicating concentrations [reviewed in (Flannagan et al., 2015; Lopez and Skaar, 2018)]. On the pathogen side, bacteria have evolved several strategies to survive excess of metal ions, both in the environment (Ducret et al., 2016; Gonzalez et al., 2018) and in contact with phagocytes (Botella et al., 2011). For instance, metal efflux transporters such as cation diffusion facilitators (CDFs) and P-type ATPases remove excess ions from the bacterial cytoplasm (Chan et al., 2010; Kolaj-Robin et al., 2015).

Recently, Zn poisoning was shown to belong to the killing strategies of macrophages (Botella et al., 2011). Inside macrophages free Zn mainly localizes to late endosomes and lysosomes, and to a smaller extent to early endosomes (Botella et al., 2011). Zn presumably enters cytoplasmic organelles either by fusion with Zn-containing endosomes, called zincosomes, or by the direct action of one or several Zn transporters located at the membrane of the organelle. For example, Zn has been observed to accumulate in phagolysosomes containing the non-pathogenic bacteria *Escherichia coli*, contributing to their killing. In addition, Zn transporters are upregulated in macrophages infected with *Mycobacterium tuberculosis* (*Mtb*) and, after a cytosolic burst, free Zn is delivered and accumulates into the *Mycobacterium*-containing vacuole (MCV) 24 hours after infection (Botella et al., 2011; Pyle et al., 2017; Wagner et al., 2005). Interestingly, vice versa, the expression of the P_1_-type Zn exporting ATPase CtpC of *Mtb*, the Zn exporting P_1B_-type ATPase ZntA of *E. coli* and *Salmonella enterica* serovar Typhimurium, and the Zn Cation Diffusion Facilitator of *Streptococcus pyogenes* increase during infection of human macrophages and neutrophils (Botella et al., 2011; Kapetanovic et al., 2016; Ong et al., 2014).

However, the import mechanism and the temporal dynamics of Zn inside phagosomes containing non-virulent and virulent bacteria are still poorly understood. Two families of Zn transporters, ZnT (Zn transporter Slc30a) and ZIP (Zrt/Irt-like protein Slc39a), have been described in metazoans and in the amoeba *Dictyostelium discoideum* (hereafter referred to as *Dictyostelium*) [reviewed in (Dunn et al., 2017; Kambe et al., 2015)]. In metazoans, the ZnT family consists of nine proteins (i.e. ZnT1-8 and ZnT10) that decrease cytosolic Zn levels by exporting it to the extracellular space or sequestering it into the lumen of organelles. In contrast, the fourteen members of the ZIP family (i.e. ZIP1-14) catalyse the transport of free Zn from the extracellular space and organelles into the cytosol [reviewed in (Kambe et al., 2015)]. In *Dictyostelium*, four putative ZnT proteins and seven ZIP transporters have been identified, but the directionality of transport has not been experimentally confirmed yet (Dunn et al., 2017; Sunaga et al., 2008).

In recent years, *Dictyostelium* has evolved as a powerful model system to study phagocytosis and host-pathogen interactions, including cell autonomous defences and bacterial killing [reviewed in (Bozzaro et al., 2008; Dunn et al., 2017)]. The core machinery of the phagocytic pathway is highly conserved between human phagocytes and *Dictyostelium*, which digests and kills non-pathogenic bacteria for nutrition. First, bacteria are recognized by receptors at the plasma membrane, leading to an actin-dependent deformation of the membrane and the closure of the phagocytic cup (Neuhaus et al., 2002). Few minutes after uptake, lysosomes fuse with the phagosome, delivering the vacuolar H^+^-ATPase (vATPase), which acidifies the phagosomal lumen (pH < 4) and contributes to the digestion of bacterial components in concert with various sets of lysosomal enzymes that are transferred to the phagosome by fusion with endosomes and lysosomes (Gotthardt et al., 2002). These enzymes are then retrieved from the phagosomes together with the vATPase starting 45 min after uptake, leading to a post-lysosomal compartment with neutral lumenal pH (Gotthardt et al., 2002). Undigested material is then expelled from the cell by exocytosis. In *Dictyostelium*, free Zn has been recently localized in the contractile vacuole (CV), an organelle crucial for osmoregulation, and in the endo-lysosomal system, including phagosomes containing non-pathogenic *E. coli* (Buracco et al., 2017).

Here, we monitor free Zn in the endo-phagocytic pathways of *Dictyostelium*, to compare to knowledge acquired in macrophages, and to understand the evolutionary origin of the metal poisoning strategy. Moreover, so far it remains unclear whether ZnTs are involved in the Zn accumulation in the endosomal-lysosomal pathway and consequently we extended our analysis to various *znt* knockouts.

## Results

### In *Dictyostelium*, zincosomes are mainly of lysosomal and post-lysosomal nature

To decipher in more detail the recently reported subcellular localization of free Zn in *Dictyostelium* (Buracco et al., 2017), cells were pre-incubated overnight with the fluid phase tracer TRITC-dextran, which accumulates in all endosomes and lysosomes (Hacker et al., 1997), stained with either Fluozin-3 AM (FZ-3, Fig. 1A) or N1-(7-Nitro-2,1,3-benzoxadiazol-4-yl)-N1,N2,N2-tris(2-pyridinylmethyl)-1,2-ethane-diamine (NBD-TPEA, Fig. 1B), two fluorescent probes selective for Zn, and fed with 3 μm latex beads. Importantly, using these two fluorescent probes we were able to localize Zn inside the different cellular compartments, but not in the cytosol.

**Fig. 1.**
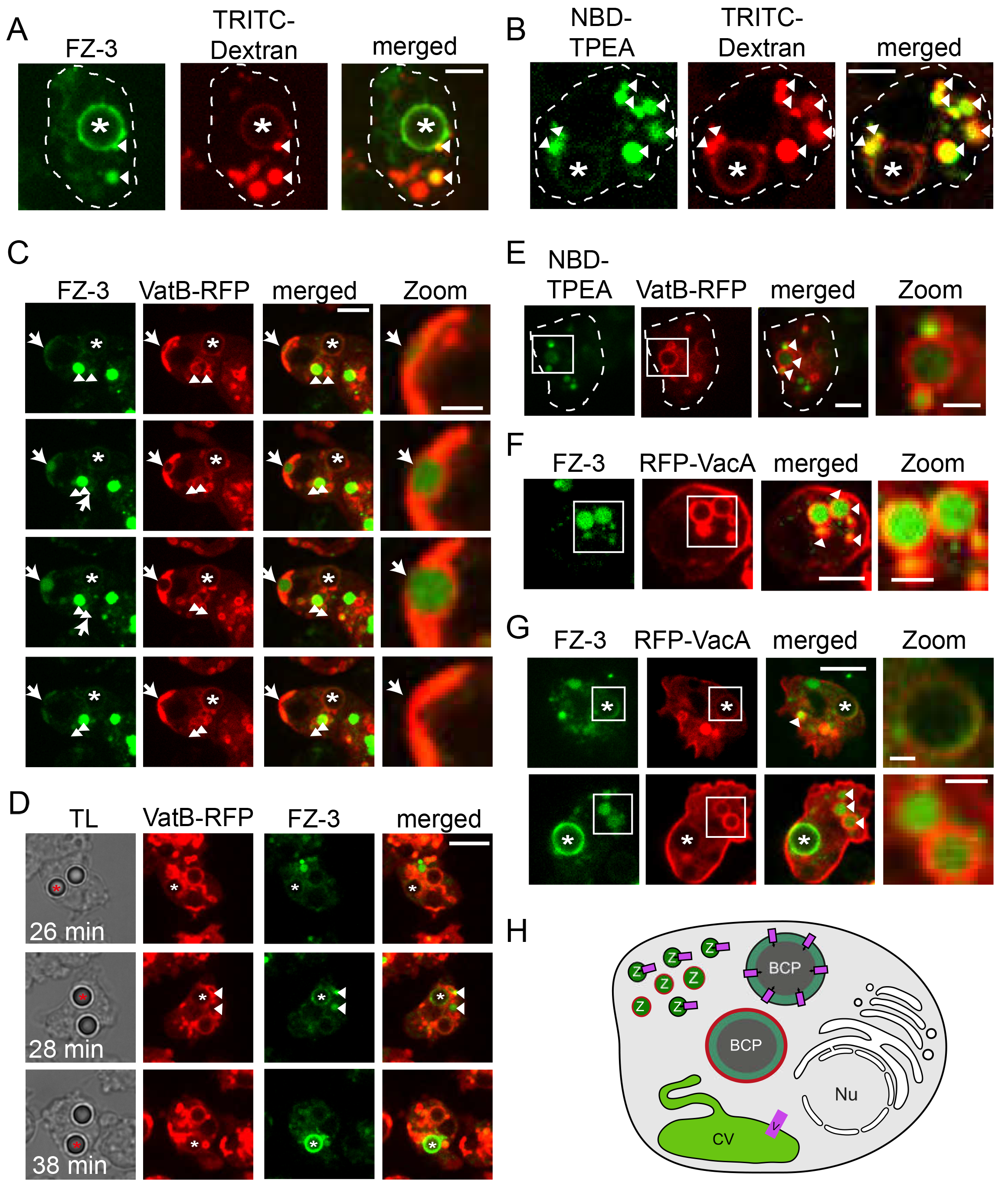
Zn accumulates inside the CV network and zincosomes. **A.-B.** Zn is observed in the endosomal system. Cells were incubated with TRITC-dextran, stained with FZ-3 (A) or NBD-TPEA (B), and fed with 3 μm latex beads. Arrowheads point to zincosomes. **C**. Zinc is removed from the cell when the CV discharges. Arrows point to the site of Zn discharge from the CV. Shown are snapshots from Movie 1. Scale bar, 5 μm; Zoom 2 μm. **D.** FZ-3 fluoresces inside a BCP when the vATPase is retrieved. Scale bar, 10 μm. TL: transmitted light. **E.** Zincosomes have partially lysosomal characteristics. Scale bars, 5 μm; Zoom 2 μm. Cells expressing VatB-RFP were stained with FZ-3 (C and D) or NDB-TPEA (E) and fed with beads. **F.** Zincosomes have partially post-lysosomal characteristics. Scale bars, 5 μm; Zoom 2 μm. **G.** Zn accumulates inside BCPs at the post-lysosomal stage. Scale bars, 5 μm; Zoom 1 μm. RFP-VacA-expressing cells were used (F and G). **H.** Scheme. Zn accumulates inside the CV network, zincosomes and the BCP in *Dictyostelium.* Asterisks label BCPs. BCP: Bead-containing phagosome, v: vATPase, CV: contractile vacuole, Nu: Nucleus, Z: zincosomes. VacA is shown in red.

As revealed by FZ-3 and NBD-TPEA staining (Fig. 1A,B), Zn was clearly detectable inside the bead-containing phagosome (BCP, asterisk) and zincosomes, that colocalised with the endosomal marker TRITC-dextran (arrowheads). To define more precisely the identity of zincosomes, *Dictyostelium* expressing VatB-RFP, a marker of lysosomes and the contractile vacuole (Bracco et al., 1997), or RFP-VacA, a marker of post-lysosomes (Wienke et al., 2006), were stained with both Zn-probes and fed with latex beads (Fig. 1C-G). Using these markers Zn was detected inside bladders of the CV before discharge (Fig. 1C, arrows, Movie 1), in VatB-RFP-positive lysosomes (Fig. 1E, arrowheads) and in VatB-RFP-negative but RFP-VacA-positive post-lysosomes (Fig. 1D,F,G; arrowheads). Besides the osmoregulatory function of the CV and its role in Ca^2+^ sequestration, it is proposed to serve as a transient sink for divalent metals (Bozzaro et al., 2013). Consequently, our observations are in line with the hypothesis that toxic metals and other ions are expelled from the cell via the CV (Bozzaro et al., 2013; Heuser et al., 1993). Importantly, we observed that lysosomes were never labelled by FZ-3 (Fig. 1D, Movie 1), whereas NBD-TPEA never fluoresced inside the CV and post-lysosomes.

Taken together, our data support the localization of Zn inside zincosomes that are of lysosomal and post-lysosomal nature, as well as inside BCPs and in the CV (Fig. 1H).

### Localization of free Zn during phagocytosis in *Dictyostelium*

We precisely monitored the dynamics of appearance and accumulation of Zn in the BCP by time-lapse microscopy (Fig. 2A-D). Strikingly, the NBD-TPEA signal became visible inside BCPs very early after uptake, peaked at 15-20 min and then disappeared until the beads were exocytosed (Fig. 2A,C; Movie 2), while the FZ-3 signal was first observed only after approximately 20 min post-uptake, peaked at 35-40 min and persisted until exocytosis (Fig. 2B,D; Movie 3). We verified that the two probes were specific for Zn, since the signals of both NBD-TPEA and FZ-3 were eliminated by TPEN [N,N,N◻,N◻-tetrakis-(2-pyridylmethyl) ethylenediamine], a cell-permeant Zn chelator, but not by the non-permeant Zn chelator DTPA (diethylenetriaminepentaacetic acid) nor the calcium and magnesium (Mg) cell-impermeant chelator EDTA (ethylenediaminetetraacetic acid, Fig. S1A-B).

**Fig. 2.**
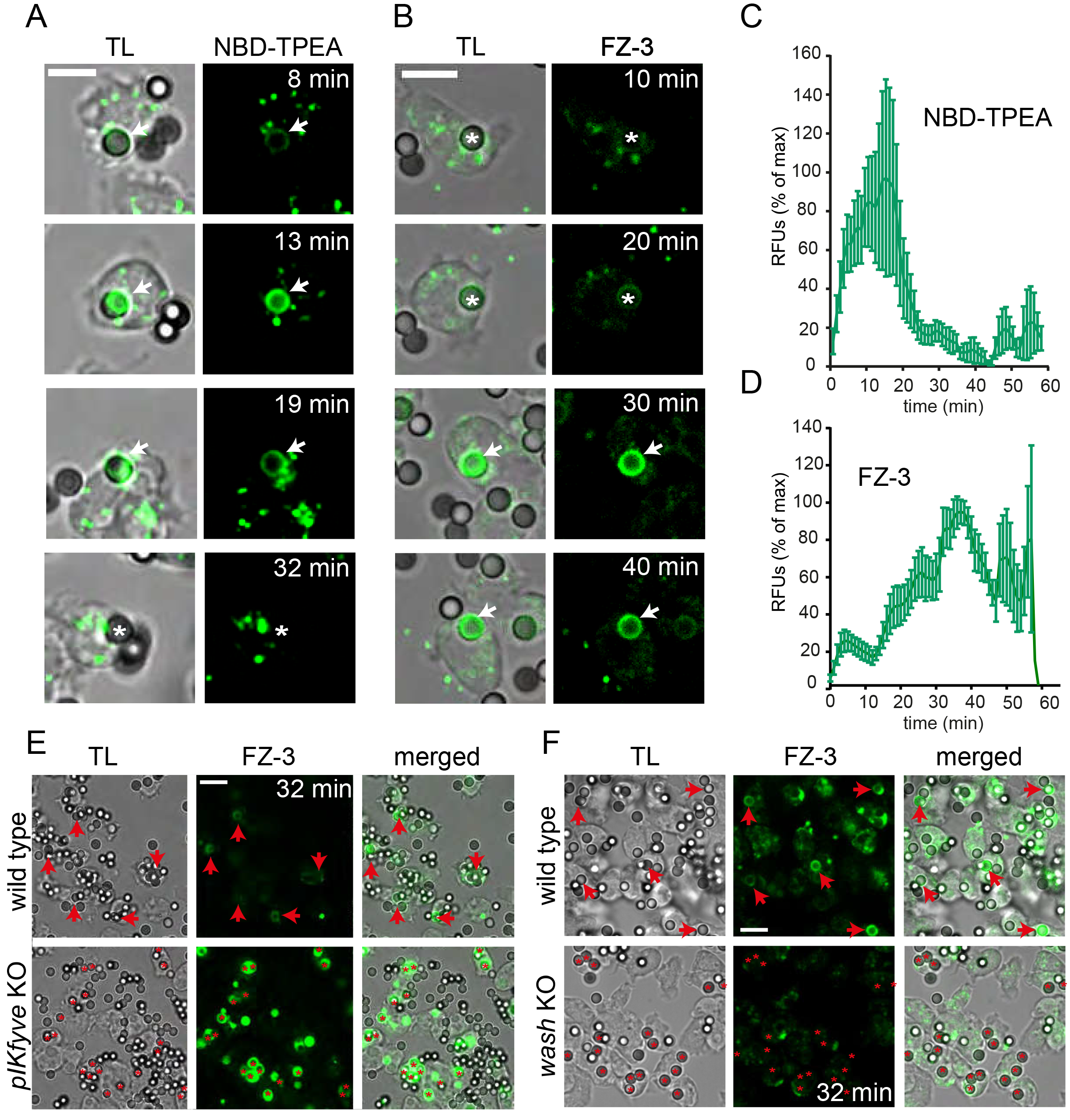
Intracellular distribution of Zn during the phagocytic pathway of *Dictyostelium.* **A.-D.** After NBD-TPEA (A) or FZ3 (C) staining, cells were fed with 3 μm latex beads. Single beads were followed by live microscopy from uptake to exocytosis (Movies 2 and 3). Scale bars, 5 μm. Relative fluorescence intensities inside the BCPs were quantified using ImageJ and the “CenteronClick” plugin (B. and D.). Eight BCPs per fluorescent probe were quantified. **E.** and **F.** *Dictyostelium* wild type and the *pikfyve* (E) or *wash* (F) KO were stained with FZ-3 and fed with latex beads. Images were taken 32 min after bead addition. Scale bars, 10 μm. Asterisks indicate the BCPs; arrows point to NBD-TPEA- or FZ-3-positive BCPs.

It is known that a pH lower than 5 quenches by 50% the FZ-3 signal (Gee et al., 2002). On the contrary, NBD-TPEA is not affected by low pH, but quenched by Cu^2+^ concentrations around 30 μM (Xu et al., 2009). The phagosomes of *Dictyostelium*, more acidic than those in macrophages (Yates et al., 2005), reach a pH < 3.5 a few minutes after particle uptake (Marchetti et al., 2009; Sattler et al., 2013), which could explain the fact that in wild type cells most of the BCPs became FZ-3- fluorescent only after approximately 30 minutes of bead uptake (Fig. 2C,D). In line with that hypothesis, the BCPs of a *pikfyve* knockout (KO), inhibited in phagosomal acidification (Buckley et al., 2018), became FZ-3-fluorescent 2 min after uptake, and remained brightly fluorescent until exocytosis (Fig. 2E). In sharp contrast, the BCPs of the *wash* mutant, which cannot reneutralize phagosomes due to a defect in the retrieval of the vATPase (Carnell et al., 2011), did not show any FZ-3-fluorescence (Fig. 2F). These results confirm that FZ-3 is quenched in early BCPs, due to the low lumenal pHs, and becomes fluorescent only during reneutralization, when the vATPase is retrieved (Fig. 1D). Regarding the sensitivity of NBD-TPEA to Cu, cells were pre-treated with different concentrations of CuSO_4_ before incubation with the probe and beads. Five μM of CuSO_4_ was sufficient to completely quench the signal of NBD-TPEA inside BCPs (Fig. S1C), indicating that its fluorescence is likely quenched by the import of Cu into maturing BCPs. Altogether, integrating the information acquired with both dyes (Fig. 1,2, Fig. S1), we conclude that free Zn is in fact present inside BCPs from early stages after bead uptake and until exocytosis of the particle. In addition, FZ-3 and NBD-TPEA can be used in combination with fluorescent organelle reporters to decipher the localization and function of Zn during grazing on bacteria.

### Zn can be delivered to phagosomes by fusion with endo-lysosomal compartments

In macrophages, Zn can be delivered to phagosomes via fusion with early endosomes (Botella et al., 2011). We wondered whether this was also the case in *Dictyostelium*. Careful examination of amoebae during the phagocytosis assays mentioned above (i.e. preincubation with TRITC-dextran, staining with FZ-3 or NBD-TPEA and feeding with 3 μm latex beads) revealed that zincosomes observed in the vicinity of the BCP (asteriks) can sometime be captured in the process of fusion with BCPs, during which they deliver their content into the BCP lumen, leading to a crescent shape that increased and spread over time (Fig. 3A, Movie 4, arrows). These observations suggest that large amounts of Zn are delivered to maturing BCPs via zincosome-phagosome fusion. Interestingly, a large fraction of the delivering zincosomes were surrounded by AmtA, an ammonium transporter present in endosomes and phagosomes (Kirsten et al., 2008; Uchikawa et al., 2011), confirming their endosomal nature (Fig. 3B).

**Fig. 3.**
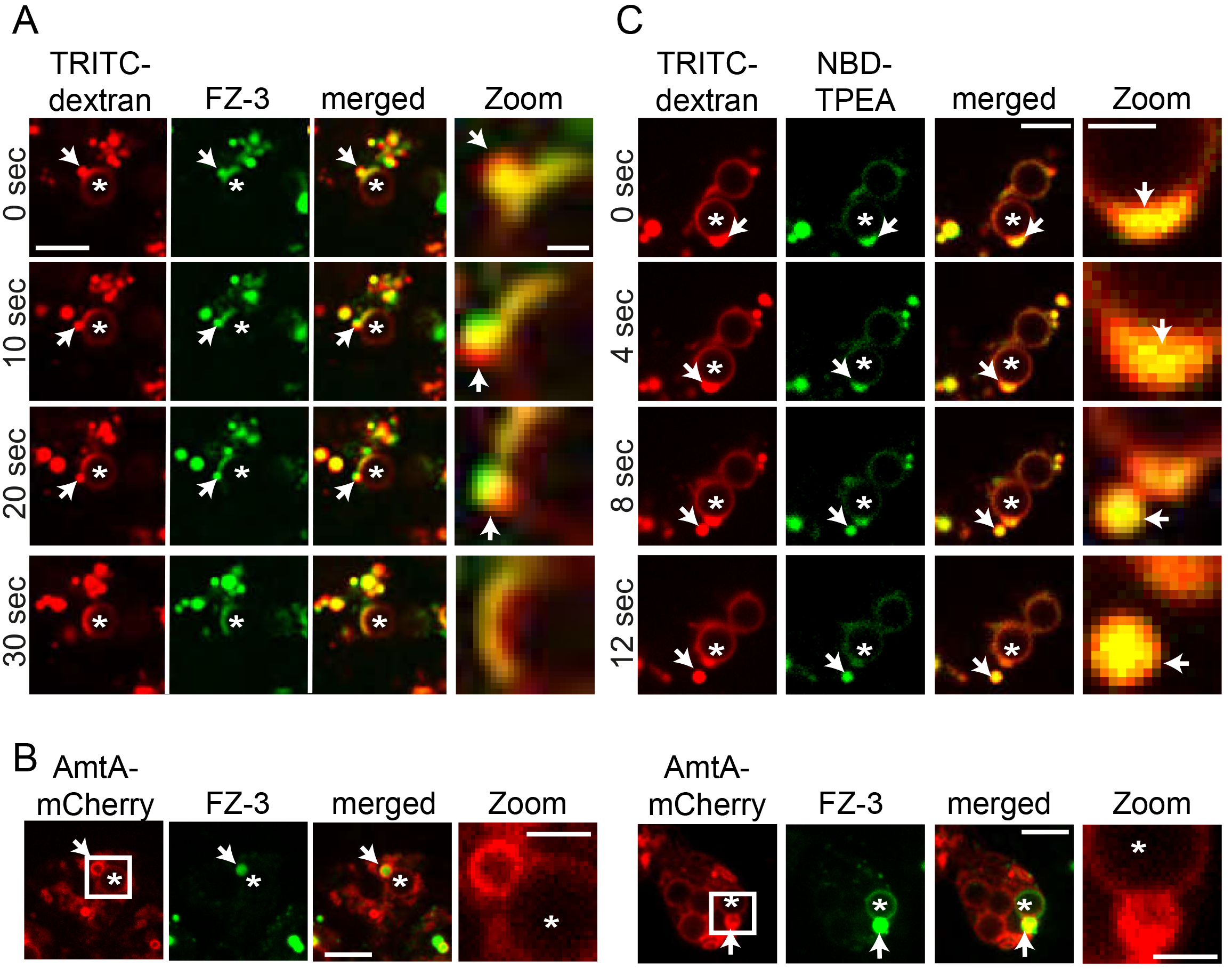
Fusion and fission dynamics of zincosomes at the BCPs. **A.** Cells were incubated with TRITC-dextran overnight, stained with FZ-3, and fed with 3 μm latex beads. Scale bar, 5 μm; Zoom 1 μm. Shown are snapshots from time-lapse Movie 4. **B.** Cells expressing AmtA-mCherry were stained with FZ-3 and fed with beads. Shown are two examples of AmtA-positive zincosomes fusing with BCPs. Scale bar, 5 μm Zoom, 1 μm. **C.** Cells were treated as in A, but stained with NBD-TPEA. Scale bar, 3 μm; Zoom 1 μm. Shown are snapshots from time-lapse Movie 5. Asterisks mark the BCPs; arrows label the zincosome-phagosome fusion site (A and B) or the zincosome-phagosome budding site (C).

On the other hand, it was also observed that Zn, detected with NBD-TPEA, was retrieved from BCPs (asterisks) by fission leading to the formation of new zincosomes (Fig. 3C, Movie 5, arrows). In summary, one way to deliver Zn to and retrieve it from phagosomes in *Dictyostelium* is by fusion and fission events with zincosomes of endosomal origin.

### Subcellular localization of ZnTs in *Dictyostelium*

Zn is sequestered into cellular organelles by Zn transporters of the ZnT family [reviewed in (Kambe et al., 2015)]. In order to determine the localization of the *Dictyostelium* ZnT transporters, cells expressing fluorescent chimeras of each of the four ZnT proteins were fixed and stained with antibodies against several CV and endosomal markers (Fig. 4 and 5). ZntA-mCherry partially co-localized with VatA, a subunit of the vATPase present in both lysosomes and membranes of the CV network (Neuhaus et al., 1998), and perfectly co-localized with Rhesus50, a protein that localizes exclusively to the CV [(Benghezal et al., 2001); Fig. 4A].

**Fig. 4.**
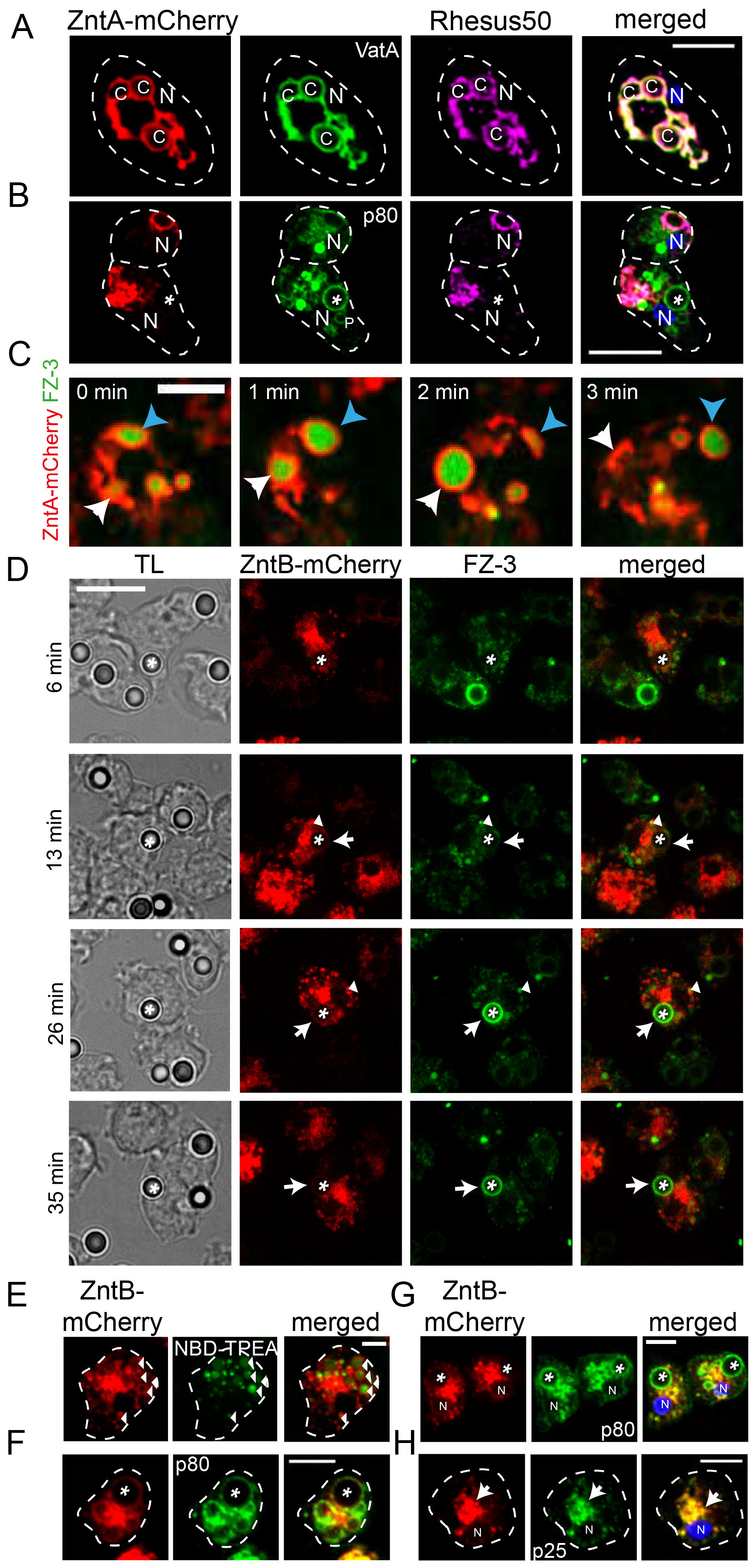
Subcellular localization of ZntA and ZntB. **A. and B.** ZntA co-localizes with the CV markers Rhesus50 and VatA, but not with the endosomal protein p80. ZntA-mCherry expressing cells were fixed and stained with antibodies against Rhesus50 (A and B), VatA (A) and p80 (B). The signal from ZntA-mCherry was enhanced with an anti-RFP antibody. Nuclei were stained with DAPI. P: post-lysosome, C: tubular network of the CV, N: nucleus. Asterisks label BCPs. Scale bars, 5 μm (A) and 10 μm (B). **C.** Zn is enriched at ZntA-labeled CV compartments. ZntA-mCherry expressing cells were stained with FZ-3. Scale bar, 5 μm. Arrows point to the sites of CV discharge. **D.** ZntB is recruited to the BCP during phagosome maturation. Shown are images from a time-lapse movie (Movie 6). Scale bar, 15 μm. **E.** ZntB localizes at zincosomes. Images were taken live. Arrowheads label zincosomes. Scale bar, 2 μm. Cells expressing ZntB-mCherry were stained with FZ-3 (D) or NBD-TPEA (E) and fed with 3 μm latex beads. Asterisks label the BCP, arrows point to a ZntB-positive BCP, arrowheads label Znt-B-mCherry-positive zincosomes. **F. - H.** ZntB co-localizes with endosomal markers. ZntB-mCherry expressing cells were fixed and stained with antibodies against p80 (F and G) and p25 (H). The signal from ZntB-mCherry was enhanced with an anti-RFP antibody. Nuclei were stained with DAPI. Asterisks label BCPs, the arrow points to a clustering of ZntB-mCherry at the juxtanuclear region. N: nucleus. Scale bars, 5 μm.

**Fig. 5.**
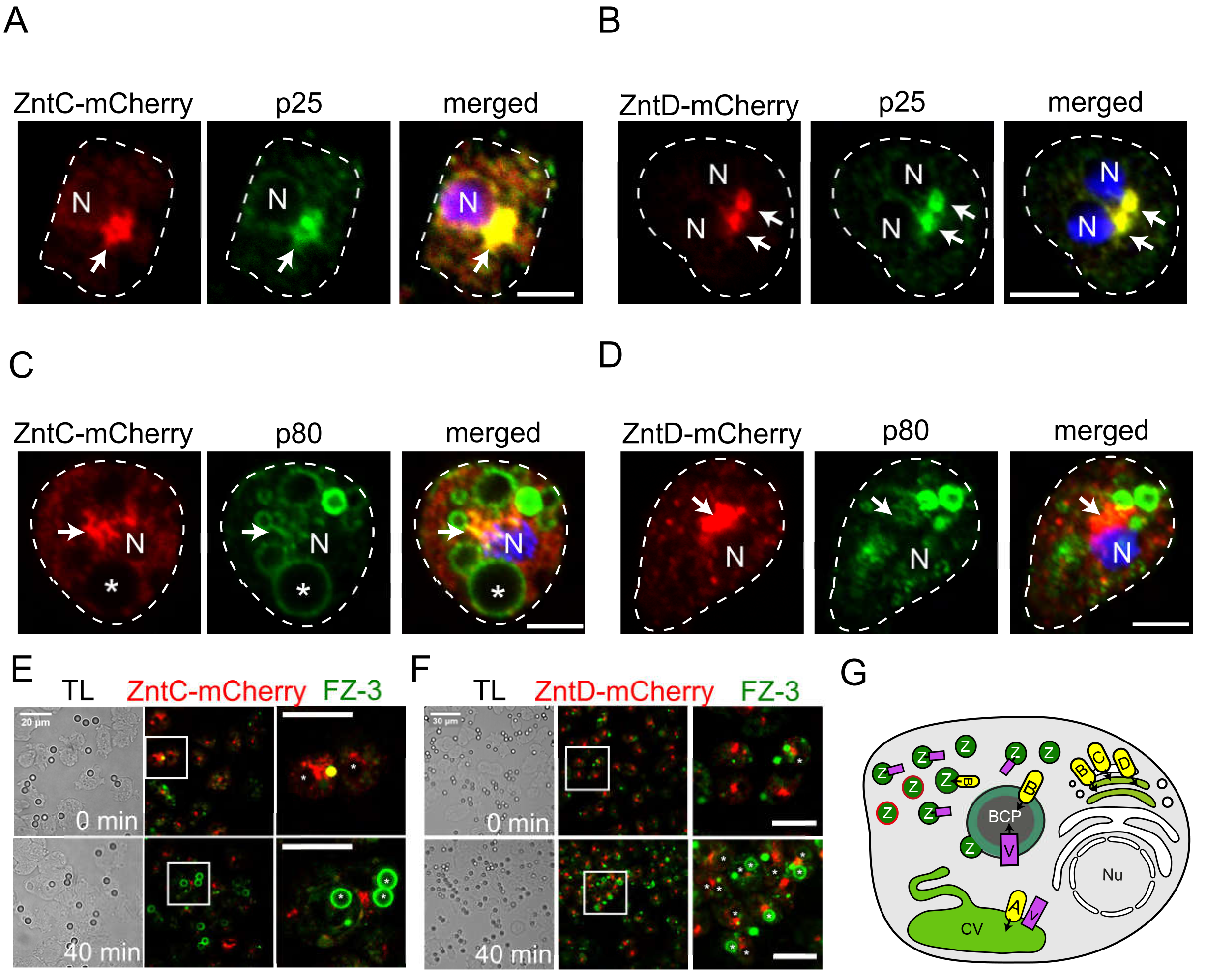
Subcellular localization of ZntC and ZntD. **A. and B.** ZntC-mCherry (A) and ZntD-mCherry (B) are localized in the juxtanuclear region. **C. and D.** ZntC-mCherry (C) and ZntD-mCherry (D) do not co-localize with the endosomal marker p80. Cells were fixed and stained with antibodies against p80 and p25. Nuclei were stained with DAPI. The signal from ZntC- and ZntD-mCherry was enhanced with an anti-RFP antibody. Asterisks label a BCP, the arrow points to an enrichment of ZntC- or ZntD-mCherry at the juxtanuclear region. N: nucleus. Scale bars, 5 μm. **E. and F.** ZntC- and ZntD-mCherry do not co-localize with BCPs. Cells were stained with FZ3 and fed with 3 μm latex beads. Shown are images from time-lapse movies. Asterisks label BCPs. Scale bars 20 μm, Zoom 15 μm. **G.** Scheme summarizing the subcellular localization of ZnTs in *Dictyostelium*. BCP: Bead-containing phagosome, A: ZntA, B: ZntB, C: ZntC, D: ZntD, v: vATPase, CV: contractile vacuole, Nu: Nucleus, Z: zincosomes. VacA is shown in red.

To monitor whether ZntA-mCherry might additionally localize to phagosomes at some stage of maturation, cells were fed with latex beads, fixed and stained with an antibody against p80 (Fig. 4B). These experiments excluded colocalisation of ZntA and p80 (Fig. 4B), and highlighted that ZntA-mCherry was in fact never detected in BCPs (Fig. S2, asterisks). To confirm the exclusive presence of ZntA at CV membranes, live microscopy was performed on ZntA-mCherry-expressing cells incubated with FZ-3. The fluorescent probe accumulated in ZntA-positive bladders of the CV that frequently discharged their content, expelling FZ-3-chelated Zn to the extracellular milieu (Fig. 4C, arrows). Therefore, we conclude that ZntA is located exclusively to the CV bladders and network.

Using the same approach, cells expressing ZntB-mCherry and incubated with FZ-3 were fed latex beads, revealing the presence of ZntB-mCherry at BCPs (Fig. 4D asterisks) starting from 6 min to until 35 min after bead uptake (arrows, Movie 6). In addition, ZntB-mCherry was also observed around FZ-3-positive zincosomes (Fig. 4D, arrowheads). The ZntB-mCherry-labelled zincosomes (Fig. 4E, arrowheads) and BCPs (Fig. S3A, asterisks) were also positive for NBD-TPEA, indicating that the ZntB-positive compartments can be acidic and likely contain a relatively low Cu concentration. In addition, ZntB-mCherry partially co-localized with p80 at BCPs (Fig. 4F,G), indicating their lysosomal or early post-lysosomal nature. This is in line with the observation that ZntB-mCherry also co-localized with the vATPase (Fig. S3B) but neither with Rhesus50 nor with the endoplasmic reticulum marker PDI (Fig. S3B). Similar to its human homolog ZNT10 (Bosomworth et al., 2012; Dunn et al., 2017), ZntB was also located at recycling endosomes and/or the Golgi apparatus, as determined by its juxtanuclear co-localization with p25 [(Charette et al., 2006); Fig. 4H]. Note that, in *Dictyostelium*, recycling endosomes concentrate around the Microtubule Organizing Center (MTOC), a region where typically the Golgi apparatus is also located. These results lead us to conclude that ZntB locates mainly to organelles of the endosomal pathway.

To decipher the subcellular localization of ZntC and ZntD, homologs of the human ZNT6 and ZNT7, respectively (Dunn et al., 2017), cells expressing ZntC- and ZntD-mCherry were fixed and stained with antibodies against p80 and p25, a marker for recycling endosomes. ZntC- and ZntD-mCherry both colocalized with p25 in the juxtanuclear region (Fig. 5A,B), and did not colocalize significantly with p80 (Fig. 5C,D), suggesting that these transporters either locate in recycling endosomes that are usually low in p80 (Charette et al., 2006), or in the Golgi apparatus. Importantly, ZntC- and ZntD-mCherry were not observed at BCPs at any stage of maturation (Fig. 5E,F). The locations of the various *Dictyostelium* ZnTs are schematized in Fig. 5G.

### ZntA is the main Zn transporter of the CV

Zn accumulates inside the CV and is expelled from the cell when the CV discharges (Fig. 1C, 4C). Since ZntA was the only ZnT localized at the CV membrane (Fig. 4A-C, Fig. S2), we wondered whether ZntA was involved in the import of Zn into the CV lumen. To test this hypothesis, a *zntA* KO was created (see Materials and Methods and Fig. S4A,B), and wild type and *zntA* KO cells expressing VatB-RFP were incubated with FZ-3 (Fig. 6A,B). To verify that the absence of ZntA did not alter the dynamics of Zn delivery into BCPs, cells were fed with 3 μm latex beads during the assay. No difference in FZ-3 signal intensity nor in Zn delivery to BCPs were observed between wild type and mutant cells (Fig. 6A-B). However, Zn, which was found in the VatB-RFP-positive CV of wild type cells at all time points (Fig. 6A), was strikingly absent from the CV of *zntA* KO cells (Fig. 6B). This suggests that ZntA mediates the main route of Zn delivery into the CV. Overexpressing mCherry-ZntA in *zntA* KO cells rescued the transport of Zn inside the CV network (Fig. 6C). Interestingly, when cells expressing AmtA-mCherry were stained with NBD-TPEA, the intensity of the signal in zincosomes of the *zntA* KO was more intense than in wild type cells (Fig. 6D, E). Importantly, knocking out *zntA* did not alter the number of zincosomes (Fig. 6F).

**Fig. 6.**
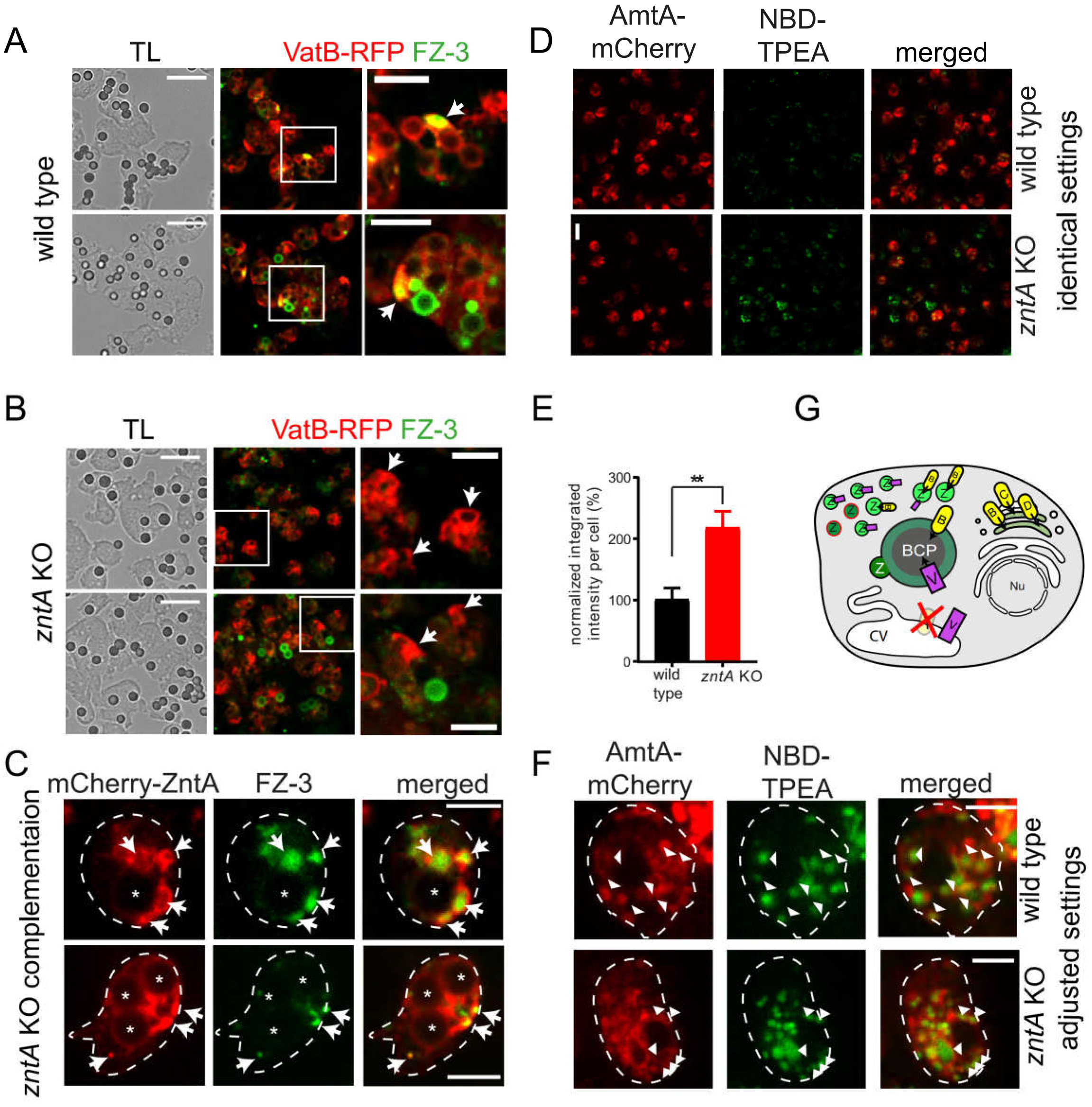
ZntA is the main Zn transporter of the CV. **A. and B.** Zn is absent from the CV network of the *zntA* KO. Shown are pictures from time-lapse movies taken at 8 min (upper panel) and 50 min (lower panel) after the experiment was started. Scale bar, 20 μm; Zoom 10 μm. **C.** Overexpression of mCherry-ZntArescues the phenotype of the *zntA* KO. Scale bars, 5 μm. Wild type (A) and *zntA* KO (B) expressing VatB-RFP or *zntA* KO cells expressing mCherry-ZntA (C) were fed with 3 μm latex beads and stained with FZ3. The arrows point to CV bladders filled with Zn in the wild type or mCherry-ZntA overexpressor and to empty CV bladders in the *zntA* KO. Asterisks label BCPs. **D.** Mislocalized Zn is shuttled into acidic zincosomes in the *zntA* KO. Cells expressing AmtA-mCherry were stained with NBD-TPEA. Images were taken by live-microscopy. Scale bar, 10 μm **E.** Quantification of D. The total integrated density per cell was quantified using ImageJ. Statistical significance was calculated with an unpaired t-test (**P < 0.01). Bars represent the mean and SEM of two independent experiments. In both conditions 30 cells were analysed. **F.** Same than D, but the microscope settings were adjusted to compare the number of zincosomes in wild type and *zntA* KO. Arrowheads point to zincosomes. Scale bars, 5 μm. G. Summarizing scheme showing the mislocalization of Zn in the *zntA* KO. BCP: Bead-containing phagosome, A: ZntA, B: ZntB, C: ZntC, D: ZntD, v: vATPase, CV: contractile vacuole, Nu: Nucleus, Z: zincosomes. VacA is shown in red.

In conclusion, these data indicate that ZntA is the main Zn transporter of the CV system, and that the absence of ZntA leads to an increased concentration of Zn in the endosomal system (Fig. 6G).

### ZntB is the main lysosomal and post-lysosomal Zn transporter

As mentioned above, ZntB localized at zincosomes and phagosomes (Fig. 4D-H, Fig. S3). To test whether ZntB mediates the transport of Zn into these compartments, a *zntB* KO, generated within the Genome Wide *Dictyostelium* Insertion (GWDI) project, was used (Fig. S4C). The insertion site of the blasticidin cassette was confirmed by genomic PCR (Fig. S4D). Wild type and *zntB* KO cells were incubated over night with TRITC-dextran, stained with FZ-3 or NBD-TPEA, and fed with 3 μm latex beads (Fig. 7A-D). Strikingly, the BCPs of *zntB* KO cells appeared to be devoid of both FZ-3 and NBD-TPEA signals (Fig. 7A and D). By adjusting the image settings and by quantification of the integrated signal intensity inside BCPs, the signal in *zntB* KO cells was shown to be decreased by approximately 60% compared to wild type (Fig. 7B and C). The residual amount of Zn detected might be delivered to BCPs by fusion with zincosomes that are also present in the *zntB* KO (Fig. 7D, arrows). Importantly, overexpression of ZntB-mCherry in the *zntB* KO rescued the defect in Zn content (Fig. 7E).

**Fig. 7.**
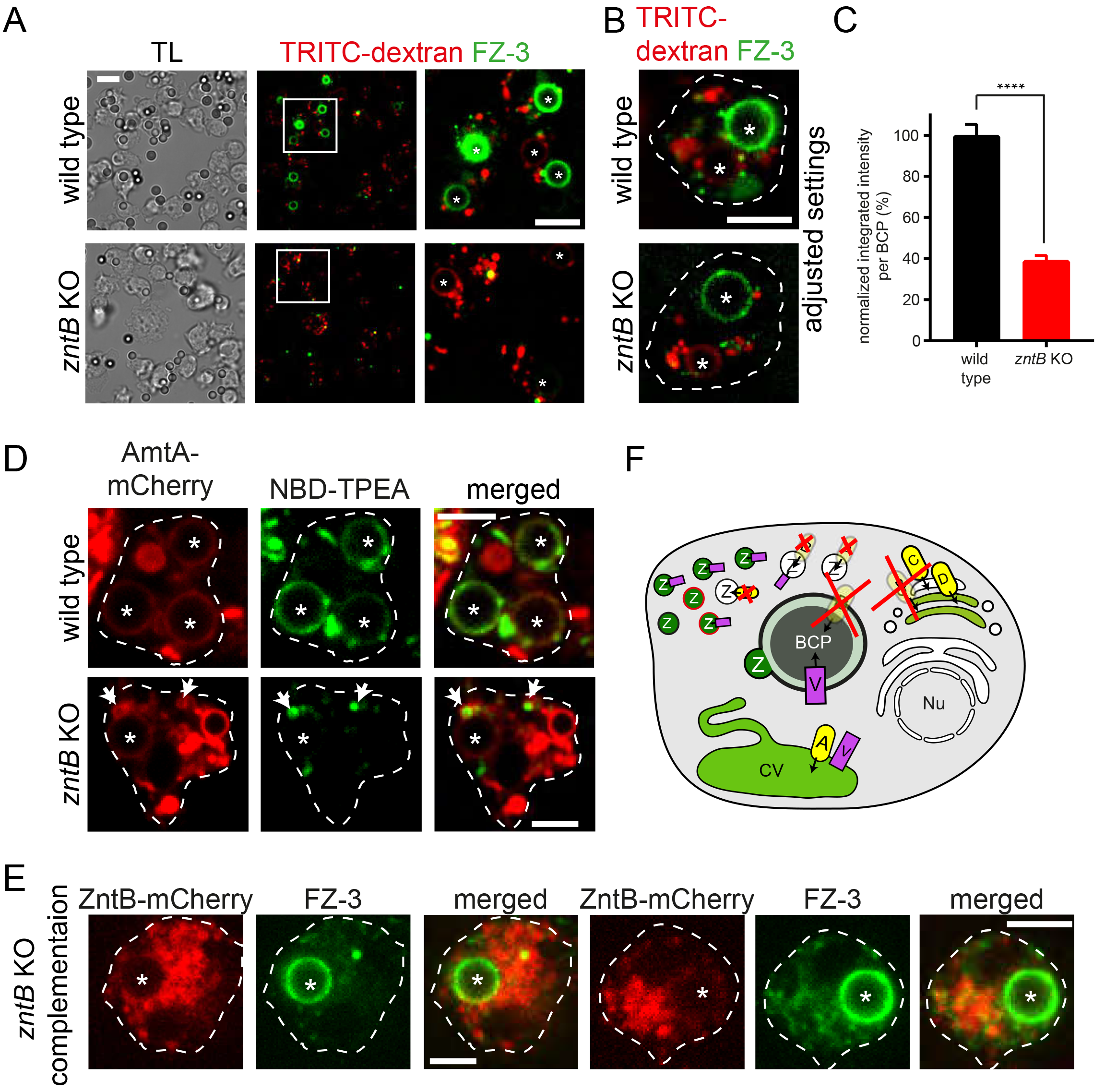
ZntB is the main endosomal Zn transporter. **A.-D.** Zn is almost absent from BCPs in the *zntB* KO. **A.** Cells were incubated over night with TRITC-dextran, co-stained with FZ-3 and fed with 3 μm latex beads. Scale bar, 10 μm; Zoom 5 μm. **B.** Same than A. To show the residual amount of Zn inside the BCP of the *zntB* KO, the brightness and contrast of the image was adjusted using ImageJ. Scale bar, 5 μm. **C.** Quantification of A. The integrated density inside the BCPs was quantified using ImageJ. Statistical significance was calculated with an unpaired t-test (****P < 0.0001). Bars represent the mean and SEM of three independent experiments. 103 BCPs were analysed for the wild type and 93 for the *zntB* KO. **D.** Cells expressing AmtA-mCherry were stained with NBD-TPEA and fed with 3 μm latex beads. Scale bars, 5 μm. **E.** The phenotype of the *zntB* KO is rescued by overexpression of ZntB-mCherry. *ZntB* KO expressing ZntB-mCherry were incubated with FZ-3 and fed with 3 μm latex beads. Scale bars, 5 μm. Arrows point to zincosomes. Asterisks label BCPs.

We propose that ZntB is the main Zn transporter in lysosomes and post-lysosomes (Fig.7F), and that, in its absence, residual levels of Zn are reached within these compartments by fusion with zincosomes or trafficking from recycling endosomes, where Zn is transported by ZntC or ZntD.

Since ZntB is located at BCPs, a generic type of phagosomes that have a relatively transient nature, leading to particle exocytosis after about 60 minutes, we wondered whether ZntB is also present at compartments containing the non-pathogenic mycobacterium *M. smegmatis*, which are documented to have phagolysosomal identity but are more persistent, releasing killed bacteria after a couple of hours. ZntB-mCherry localized at the membrane of the MCV immediately after bacteria uptake, and Zn, detected by NBD-TPEA, also accumulated inside the MCV at the same time (Fig. S5). The concentration of intraphagosomal Zn appeared to increase during the early stages of infection (from 1 to 33 min post-uptake), which agrees with the hypothesis of ZntB being the main lysosomal Zn transporter. Strikingly, as we observed before for BCPs (Fig. 7D), Zn could also be delivered to the MCV by fusion of ZntB-mCherry-decorated zincosomes (Fig. S5, 33 min).

### Zn poisoning contributes to the killing of phagocytosed bacteria

In macrophages, infection with *E. coli* leads to an increase in the cytosolic level of Zn, followed by its transport inside phagosomes (Botella et al., 2011). When *Dictyostelium* was infected with *E. coli*, we could not observe a cytosolic burst of Zn, but Zn appeared in the phagosomes rapidly after uptake (Fig. 8A). This was due, at least in part, to fusion of the phagosomes with zincosomes (Fig. 8A, arrowheads) in line with our previous observations (Fig. 3A,S5) and the findings in macrophages (Botella et al., 2011).

**Fig. 8.**
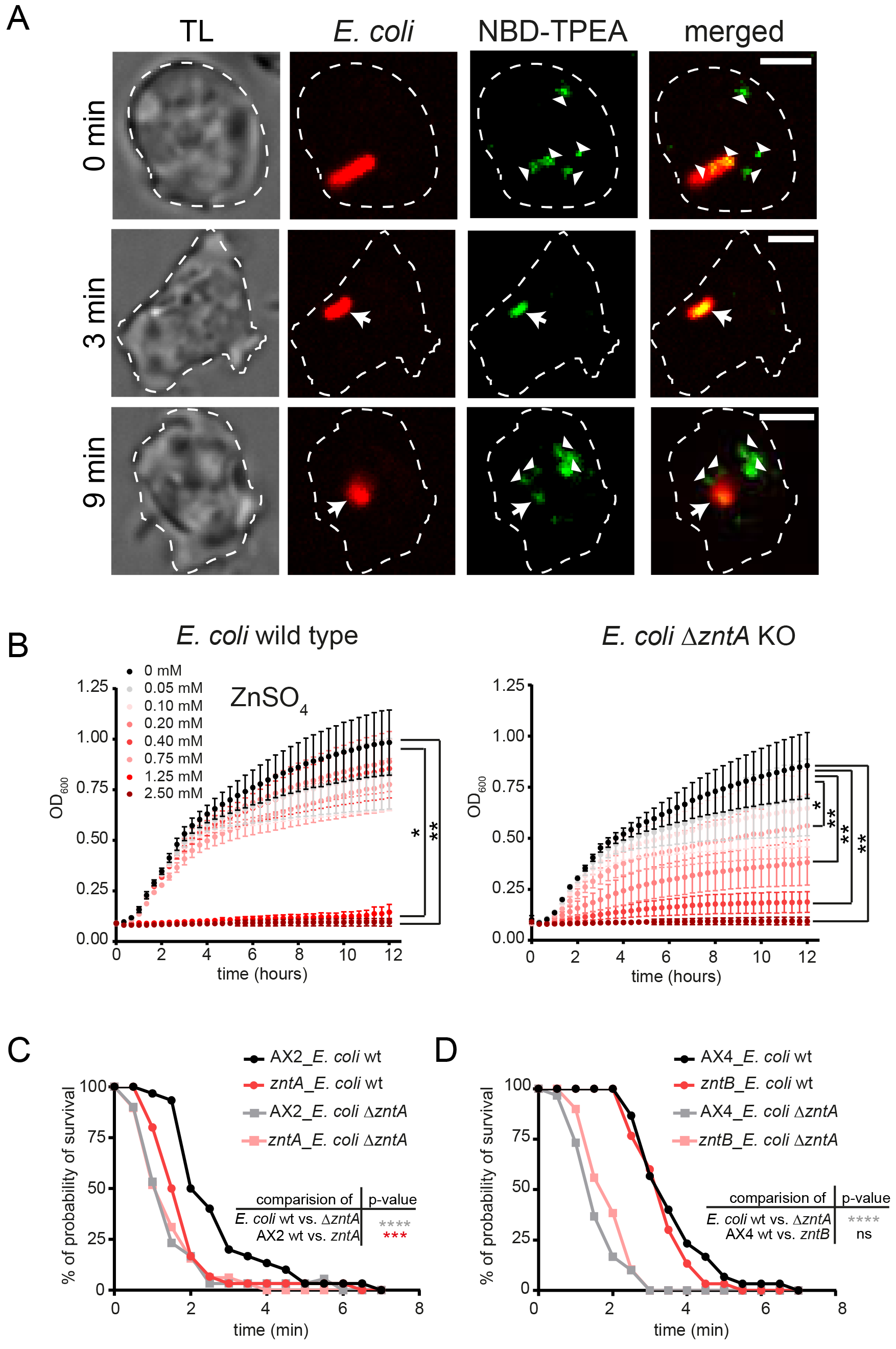
Zn poisoning is a killing strategy of *Dictyostelium.* **A.** Zn accumulates in *E. coli*-containing phagosomes. Cells were incubated with NBD-TPEA and fed with CF594-labelled bacteria before live-imaging. Scale bars, 5 μm. Arrows point to phagosomes, arrowheads label zincosomes. **B.** An *E. coli ΔzntA* mutant is more sensitive to increasing concentration of ZnSO_4_ than the wild type. *E. coli* stains were incubated in LB. ZnSO_4_ was added as indicated. The OD_600_ was measured with a 96-well plate reader (SpectraMax i3, Molecular Devices). Statistical differences are calculated with a Bonferroni post hoc test after two-way ANOVA. Significantly different values were indicated by an asterisk (* P < 0.5, ** P < 0.01). **C.** Bacteria are killed more efficiently by the *Dictyostelium zntA* KO. **D.** Knockout of ZntB does not affect bacteria killing. *Dictyostelium* was added to wild type and *ΔzntA E. coli* immobilised on an imaging slide with poly-L-Lysine and a time-lapse movie was recorded with 30 sec intervals. For the Kaplan-Meier survival curves, the data of three independent experiments were combined. Thirty ingested bacteria were monitored per condition. The statistical significance was calculated with a log-rank test (*** P = 0.0008, **** P < 0.0001).

To determine whether Zn contributes to intraphagosomal bacteria killing by *Dictyostelium*, a Zn-hypersensitive *E. coli* mutant with an inactivated P_1B_-type Zn efflux ATPase [Δ*zntA*, (Rensing et al., 1997)] was used. We first confirmed the hypersensitivity of this mutant to Zn (Fig. 8B), since 0.2 mM of Zn was enough to strongly inhibit its growth *in vitro*, whereas the proliferation of wild type cells was only inhibited by concentrations of Zn above 1.25 mM (Fig. 8B). This differential and dose-dependent inhibition of bacteria growth was not observed for other metals such as Cu, Fe or Mn (Fig. S6A-C). In line with observations in macrophages (Botella et al., 2011), *Dictyostelium* killed the Δ*zntA* mutant bacteria faster than the wild type *E. coli* (Fig. 8C,D). Interestingly, both *E. coli* strains were killed faster by the *Dictyostelium zntA* KO than by wild type *Dictyostelium* (Fig. 8C), while no significant difference in killing was observed between *Dictyostelium* wild type and *zntB* KO cells (Fig. 8D). This suggests that accumulation of Zn in the phagolysosomes contributes to bacterial killing and that these compartments harbour a higher Zn concentration in the *zntA* KO than in wild type *Dictyostelium*.

## Discussion

Intracellular bacteria and pathogens are limited to nutrients that are available inside the host cell. Professional phagocytes are able to exploit this dependence and have developed strategies to restrict intracellular bacteria growth or killing. For instance, essential nutrients such as Fe and Zn are either withheld from the pathogen-containing vacuole or pumped into the phagosomal lumen to ensure bacteria killing in concert with other immunity factors. Nramp1 mediates Fe sequestration from the phagosome and leads consequently to the starving of the phagosomal bacteria (Bozzaro et al., 2013; Peracino et al., 2006). Surprisingly, both nutrient deprivation (Djoko et al., 2015; Kehl-Fie and Skaar, 2010; Subramanian Vignesh et al., 2013) and metal poisoning have been reported in the case of Zn (Botella et al., 2011; McDevitt et al., 2011; Soldati and Neyrolles, 2012). Here, we investigated the subcellular localization and role of Zn in phagocytosis and killing using *Dictyostelium* as a model professional phagocyte.

Inside cells Zn is either bound to proteins or sequestered as free zinc into the various cellular compartments or vesicles. In mammals, 50% of the total cellular Zn is present in the cytoplasm, 30 to 40% is in the nucleus, and 10% is located at the plasma membrane [reviewed in (Kambe et al., 2015)]. The cytosolic Zn concentration ranges from picomolar to low nanomolar (Vinkenborg et al., 2009).

By using two fluorescent probes, FZ-3 and NBD-TPEA, we investigated the cellular distribution of compartmentalized free Zn in the professional phagocyte *Dictyostelium*. In line with previous observations from Buracco and colleagues (Buracco et al., 2017), Zn located inside the endolysosomal system and, more precisely, inside zincosomes of lysosomal and post-lysosomal nature (Fig. 1E - G and summarizing scheme Fig. 1H), as well as inside the CV network (Fig. 1C and Fig. 4C). In addition, Zn was present in the lumen of phagosomes soon after bead uptake and until exocytosis (Fig. 2A-D).

The total cellular Zn concentration varies in the range of 10-100 micromolar in mammalian cells (Krezel and Maret, 2006). It was suggested that the concentration of cytosolic Zn fluctuates in response to various stimuli (Kambe et al., 2015). For instance, after incubation of macrophages with *E. coli*, Zn is released from storage complexes, followed by pumping and sequestration into phagosomes. However, this was not observed in *Dictyostelium* neither by feeding cells with beads nor with bacteria (Fig. 2A, 8A, S5). We reason that, since the concentration of cytosolic Zn is very low [between pico- and low nanomolar; reviewed (Kambe et al., 2015)], it is consequently under the detection limit of FZ-3 and NBD-TPEA, explaining why we could not monitor cytosolic Zn.

The subcellular homeostasis of Zn is tightly regulated through uptake, storage, re-distribution and efflux mechanisms that are, among other, mediated by Zn transporters of the ZnT and ZIP family (Bird, 2015). Seven members of the ZIP family and four members of the ZnT family have been identified in *Dictyostelium* (Dunn et al., 2017; Sunaga et al., 2008). ZntC and ZntD are the *Dictyostelium* homologs of the human ZnT6 (Huang et al., 2002) and ZnT7 (Kirschke and Huang, 2003), respectively. As their mammalian counterparts (Kambe et al., 2015), they localized at recycling endosomes and/or in the Golgi apparatus of *Dictyostelium* (Fig. 5A-F, summarizing scheme Fig. 5G). ZntA, which is not closely related to any specific human Zn transporter (Dunn et al., 2017), was located at the membrane of the CV (Fig. 4A-C, Fig. S2A, summarizing scheme Fig. 5G). *Dictyostelium* ZntB is the closest homolog of human ZnT1 and ZnT10 (Dunn et al., 2017). Whereas ZnT1 is a plasma membrane protein (Palmiter and Findley, 1995), ZnT10 locates at the Golgi apparatus and at early and/or recycling endosomes (Patrushev et al., 2012), a location similar to the one of ZntB, described here. (Fig.4H). In addition, ZntB was observed at BCPs at the phagolysosomal stage (Fig. 4D-G; Fig. S3A,B). Loss of ZntB lead to a 60%-reduction of the FZ-3 fluorescence inside BCPs (Fig. 7A-E), leading to the conclusion that ZntB is the main endo-lysosomal Zn transporter in *Dictyostelium* (summarizing scheme Fig. 7F).

Interestingly, *Dictyostelium* growth was only inhibited by Zn or Cu concentrations at 50- or 500-fold the physiological levels, suggesting a very efficient control of Zn and Cu homeostasis (Buracco et al., 2017). In *Dictyostelium* cells, a classical metallothionein activity is not detected (Burlando et al., 2002) and consequently the CV was proposed to serve as a detoxification system for metal ions such as Fe, Zn and Cu (Bozzaro et al., 2013; Buracco et al., 2017; Peracino et al., 2013).

Changes in the cytosolic Zn levels under non-steady-state conditions are remedied in a process described as “muffling” by diverse mechanisms such as the cytosolic buffering by metallothioneins, the extrusion of Zn from the cell, and the sequestration of Zn into organelles (Colvin et al., 2010). In *Dictyostelium*, Zn is sequestered into the CV thanks to the transport by the ZntA (Fig. 6A,B). Because we observed an accumulation of Zn within lysosomes in *zntA* KO cells (Fig. 6A-F, summarizing scheme Fig. 6G), we conclude that, in this mutant, Zn is rerouted from the cytosol to lysosomes in order to avoid toxic levels of cytosolic Zn.

Botella and colleagues revealed metal poisoning of ingested microbes as a novel killing strategy of macrophages (Botella et al., 2011). Besides zinc poisoning one can speculate that bacteria also have to face Cu poisoning inside the phagosome. In line with that hypothesis, NBD-TPEA was quenched at later phagocytic stages (Fig. 2A,B and Fig. S1C), a plausible sign of Cu accumulation. Interestingly, the expression of the *Dictyostelium* homolog of the phagosomal copper transporter ATP7A is upregulated upon feeding with bacteria, consistent with a possible role of Cu poisoning during phagocytosis (Hao et al., 2016).

Excess Zn levels might inhibit bacterial ATP production by impairing the activity of cytochromes (Beard et al., 1995). Additionally, excess Zn might replace other metals in the active site of various enzymes or occupy non-specific binding sites (Nies, 1999). Here, we investigated the role of Zn during phagocytosis and killing of bacteria by *Dictyostelium*. Zn was observed inside phagosomes containing *E. coli* (Fig. 8A) and *M. smegmatis* (Fig. S5). Similar to the situation described in macrophages, an *E. coli* strain deficient in the Zn efflux P_1B_-type ATPase ZntA was killed faster than the wild type (Fig. 8C,D), leading to the conclusion that Zn poisoning belongs to the killing repertoire of *Dictyostelium*. While the accumulation of Zn inside lysosomes of the *zntA* KO leads to a better killing capacity of *Dictyostelium* (Fig. 8C), bacteria killing in the *zntB* KO was unaffected (Fig. 8D). This suggests that Zn poisoning is an evolutionarily conserved process and might act in concert with other killing factors such as phagosomal acidification, ROS production, and deprivation or poisoning by other metals, which would compensate for the loss of ZntB.

## Materials and Methods

### Dictyostelium plasmids, strains and cell culture

All the *Dictyostelium* material used for this article is listed below (Table 1). *Dictyostelium* Ax2(Ka) and AX4 cells were cultured axenically at 22°C in HL5-C medium (Foremedium) supplemented with 100 U/mL penicillin and 100 μg/mL streptomycin to avoid contamination. Cell lines expressing fluorescent reporters and KO cell lines were cultured in the presence of selective antibiotics [hygromycin (50 μg/ml), neomycin (5 μg/ml) or blasticidin (5 μg/ml)]. To monitor the localization of ZnTs, *Dictyostelium* was transformed with plasmids carrying the ZntA-, ZntB-, ZntC- or ZntD-mCherry constructs [pDM1044 backbone (Veltman et al., 2009)]. The *zntA* KO was generated in the Ax2(Ka) background by homologous recombination following the one-step cloning protocol previously described (Wiegand et al., 2011). In brief, left and right arms of *zntA* were amplified using the primers 5’-AGCGCGTCTCCAATGCTGCAGGGAAGTGAGGGTGTG (forward) and 5’-AGCGCGTCTCCGTTGGTTTATGTTCGTGTTCATG (reverse) and 5’-AGCGCGTCTCCCTTCCAACAATAGATCCCGAAG (forward) and 5‘-AGCGCGTCTCCTCCCCTGCAGGTGGATGTGCACTTC-5’ (reverse), and cloned into the StarGate^®^ Acceptor Vector pKOS-IBA-Dicty1 using the StarGate cloning kit. The resulting plasmid was transformed into *Dictyostelium* by electroporation, and positive clones were selected with blasticidin (Fig. S4A). Correct integration into the genome was tested by PCR using different combinations of primers: *zntA* flanking forward 5’-CGATTTGTTGTTACCTAAATATTCGTG and *zntA* flanking reverse 5’-CACCCAATTTACACTAGTTTC ACC, *zntA* inside forward 5’-GTGGTGAAGATGGTAGTAGTAGTG and *zntA* inside reverse 5’-CATGAGTACACCTAAACTTTCACG, Bsr forward 5-AGATCTTGTTGAGAAATGTTAAATTGATC and Bsr reverse 5’-TTGAAGAACTCATTCCACTCAAATATAC (Fig. S4B).

### Verification of the zntB REMI KO

The *zntB* KO (AX4 background) was obtained as part of the Genome Wide *Dictyostelium* Insertion (GWDI) Project (https://remi-seq.org/) and was generously provided by Prof Christopher Thompson. The individual mutant was obtained from the grid. To confirm the insertion site of the blasticidin cassette into the *zntB* gene, gDNA was isolated from wild type and *zntB* KO using the High Pure PCR Template Preparation Kit (Roche) and a diagnostic PCR was performed according to the recommendations on the GWDI website. Primer combinations with the two *zntB* specific primers *zntB* forward 5’- GGCAATTCCACGTTTCATCAG and *zntB* reverse 5’- GTAACGAATT GAATCCAAATCG binding approximately 400 bp up- or downstream the insertion sides and the two primers specific for the blasticidin cassette pGWDI1 5’-GTTGAGAAATGTTAAATTGATCC and pGWDI2 5’-AT AGAAATGAATGGCAAGTTAG were used to confirm the insertion.

### E. coli and M. smegmatis strains and culture

*E. coli* wild type and Δ*zntA* were kindly provided by Prof Christopher Rensing [(Rensing et al., 1997); Chinese Academy of Sciences Beijing (China)], and cultured in LB medium. *M. smegmatis* (Hagedorn and Soldati, 2007) was cultured in 7H9 medium supplemented with 10% OADC, 0.05% Tween80 and 0.2% glycerol at 32°C in shaking. Erlenmeyer flasks containing 5 mm glass beads were used to minimize clumping of bacteria. Vybrant DyeCycle Ruby Stain (Thermo Fisher Scientific) was used to stain intracellular *M. smegmatis* before live imaging was performed.

### Imaging of free Zn in Dictyostelium

The day before imaging, *Dictyostelium* was plated on 2- or 4-well ibidi dishes. Three hours before imaging, HL5-C was changed to SIH [Formedium, full synthetic medium with low Zn concentration (2.3 mg/l ZnSO_4_)]. After 2 hrs in SIH, cells were washed thrice in Soerensen buffer and stained with 2 μM FZ-3 [Thermo Fisher Scientific (#F-24195), 400 μM stock in DMSO] or 2.5 μg/ml NBD-TPEA [Sigma (#N1040), 0.5 mg/ml stock in DMSO] for 30 min in the dark. In order to synchronize phagocytosis, cells were cooled on a cold metal plate for 10 min before adding the 3 μm latex-beads [Sigma-Aldrich (#LB30)]. Beads were mixed with Soerensen buffer and added to the cells. After centrifugation at 500g for 2 min at 4°C, the medium was carefully aspirated from the dish and an agarose overlay was placed on top of the cells, as described before (Barisch et al., 2015). Cells were imaged on an inverted 3i Marianas spinning disc confocal microscope using the 63 × glycerol or 100 × oil objectives. Where indicated, cells were treated for 3 hrs with different chelators: TPEN [Sigma-Aldrich (#87641)], DTPA [Sigma-Aldrich (#D6518)], EDTA [Sigma-Aldrich (#E6758)] or CuSO_4_ × 5 H_2_O [Sigma (#C3036)] before and throughout staining with the Zn probes. To label all endosomes TRITC-dextran [70 kDa, Sigma (#T1162)] was added overnight (1 mg/ml; stock 10 mg/ml in ddH_2_O) and throughout FZ-3 and NBD-TPEA staining.

The temporal and special dynamics of Zn inside BCPs was quantified using ImageJ and the “CenterOnClick” plugin (Nicolas Roggli, University of Geneva, unpublished) that automatically centers the “clicked” particle of interest in a recalculated image for further visualization and analysis. The integrated density inside a “donut” that was drawn around the FZ-3 signal was measured using “plot Z-axis profile”.

### Antibodies and immunofluorescence

Antibodies against p80 (Ravanel et al., 2001) were purchased from the Geneva antibody platform (University of Geneva, Switzerland). An anti-RFP-antibody (Chromotek) was used to increase the fluorescence of mCherry-expressing fusion proteins. As secondary antibodies, goat anti-mouse, anti-rabbit and anti-rat IgG coupled to Alexa488, Alexa546 (Thermo Fisher Scientific) or CF640R (Biotium) were used. For immunofluorescence, *Dictyostelium* cells were fixed with cold MeOH or 4% paraformaldehyde (PFA), as described previously (Hagedorn et al., 2006). Images were recorded with a Zeiss LSM700 confocal microscope using a 63 × 1.4 NA or a 100 × 1.4 NA oil-immersion objective.

### In vitro effects of heavy metals on E. coli growth

*E. coli* wild type and Δ*zntA* were grown in LB medium overnight at 37°C in shaking at 150 rpm. Bacteria were diluted to an OD_600_ of 0.1 and plated in 96-well plates containing LB medium with different concentrations of heavy metals (i.e. ZnSO_4_, CuSO_4_, FeCl_3_, MnCl_2_ from 0.05 to 2.5 mM). The OD_600_ was measured every hour using a 96-well plate reader (SpectraMax i3, Molecular Devices).

### Killing of bacteria and involvement of Zn

Intracellular killing of *E. coli* wild type and Δ*zntA* carrying a GFP-harbouring plasmid (Valdivia and Falkow, 1997) was monitored as described previously (Leiba et al., 2017). A 1:10 dilution of overnight *E. coli* cultures was centrifuged for 4 min at 18000*g* and bacteria were resuspended in 300 μl of filtered HL5-C. 10 μl of the bacteria suspension were plated on each well of a 4-well ibidi slide and centrifuged for 10 min at 1500 rpm. 300 μl of a 1 × 10^6^ cells/ml *Dictyostelium* culture in LoFlo (synthetic low-fluorescent medium, Foremedium) were overlayed on the bacteria, and images were recorded at 22°C with a wide field microscope using the 40 × dry objective and 30 sec intervals. For each phagocytosed bacterium, the time between phagocytosis and fluorescence extinction (killing) was determined manually using the ImageJ software, and the probability of bacterial survival was represented as a Kaplan-Meyer estimator. The data of three independent experiments were pooled and statistical comparisons between Kaplan-Meier curves were calculated using the log-rank test.

To monitor the involvement of Zn in the killing of *E. coli*, *Dictyostelium* cells were stained with NBD-TPEA, as mentioned above, and bacteria were labelled using CF594 succinimidyl ester [SE, Sigma-Aldrich (#SCJ4600031)]. Briefly, an overnight culture of bacteria was diluted 1:10 in Soerensen buffer and incubated with 2 μl of a 10mM CF594 SE stock solution (in DMSO) for one hour in the dark. After two washes with Soerensen buffer, bacteria were resuspended in 1 ml of filtered HL5-C. 10 μl of the bacteria suspension were added to the pre-cooled cells on an 8-well ibidi slide and centrifuged onto cells for 1 min at 500*g* at 4°C. Images were taken with 90 sec intervals at a spinning disc confocal microscope using the 63 × objective.

## Acknowledgments

We gratefully acknowledge the imaging platform of the University of Geneva for their expert and friendly support. We thank the Genome Wide *Dictyostelium* Insertion Project (https://remi-seq.org/) for the *zntB* REMI cell line. We are grateful to Prof Michael Rensing for sharing the Δ*zntA E. coli* knockout and Dr Ana T. López-Jiménez for constructing the RFP-VacA expression vector. We thank especially Dr Elena Cardenal-Muñoz for careful reading and editing of the manuscript and thoughtful suggestions and Dr Olivier Schaad for helping with the statistics. We thank Dr Monica Hagedorn and Prof Markus Maniak for giving comments on the manuscript and Dr Olivier Neyrolles for inspiring the project. The Soldati laboratory is supported by multiple grants from the Swiss National Science Foundation. Thierry Soldati is a member of iGE3 (www.ige3.unige.ch).

## Supporting Information

**Fig. S1.**
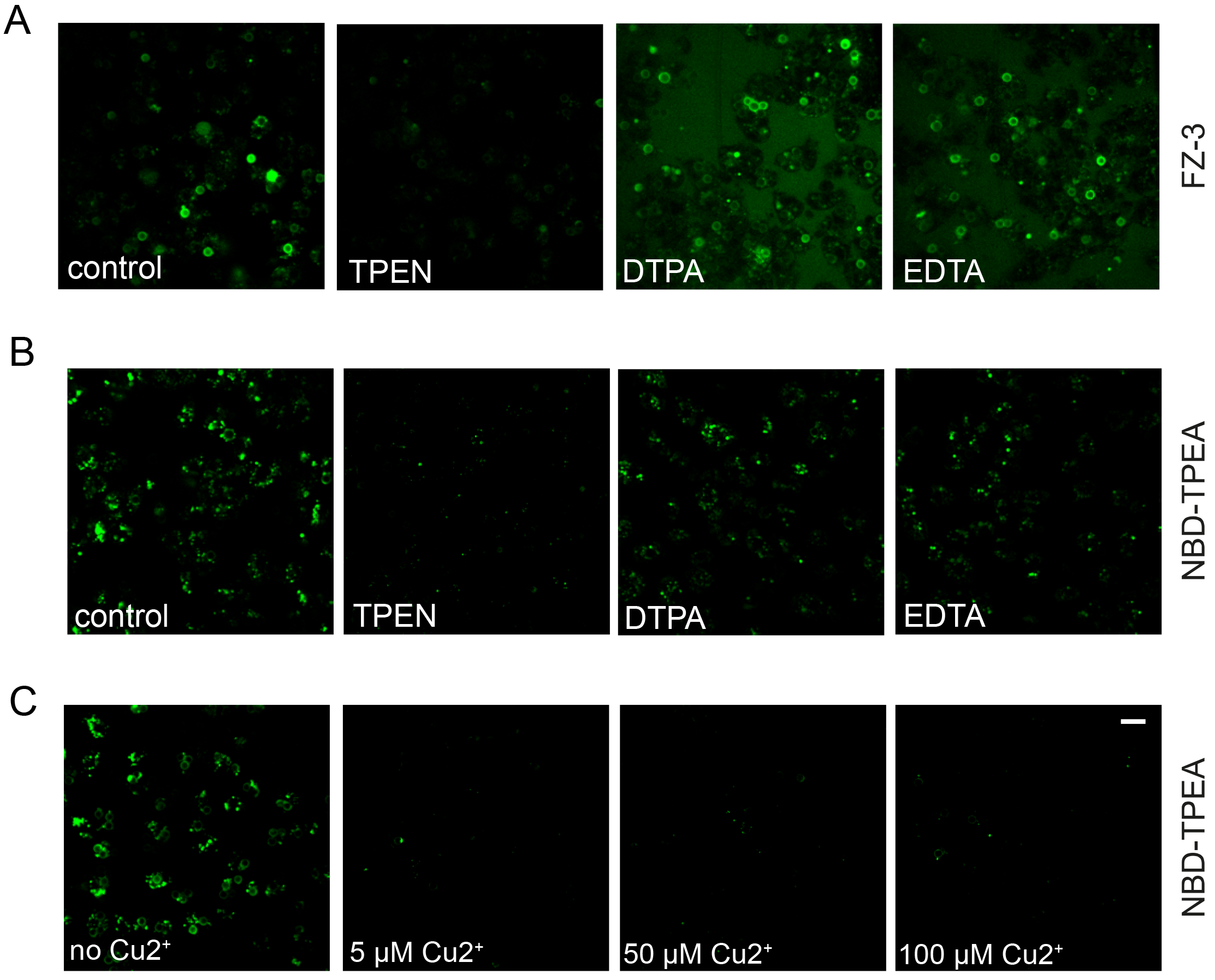
Control experiments showing the specificity of FZ-3 and NBD-TPEA for Zn and the quenching of NBD-TPEA by Cu. **A.-C.** Cells were incubated with the different chelators or CuSO_4_ at the indicated concentrations for 3 hrs before the staining with FZ-3 and NBD-TPEA was performed. Afterwards cells were fed with 3 μm and images were taken by live-microscopy. Importantly, the chelators and CuSO_4_ were present throughout the experiment.

**Fig. S2.**
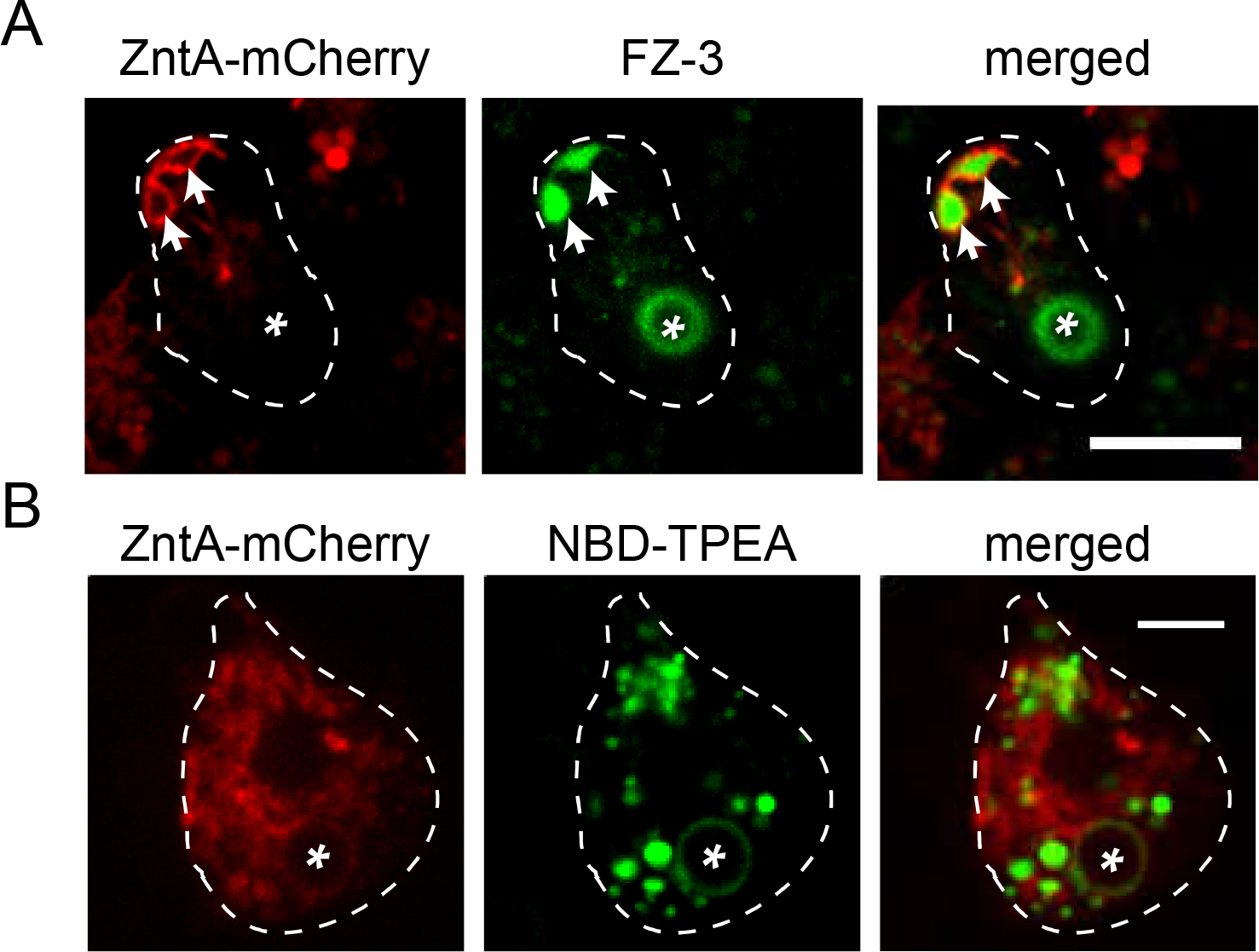
ZntA-mCherry is not present on BCPs. **A.** and **B.** Cells expressing ZntA-mCherry were stained with FZ-3 and NBD-TPEA and fed with 3 μm latex beads. Images were taken live. Asterisks label BCPs, arrows point to FZ-3-positive CV-bladders. Scale bars 10 μm (A), 5 μm (B).

**Fig. S3.**
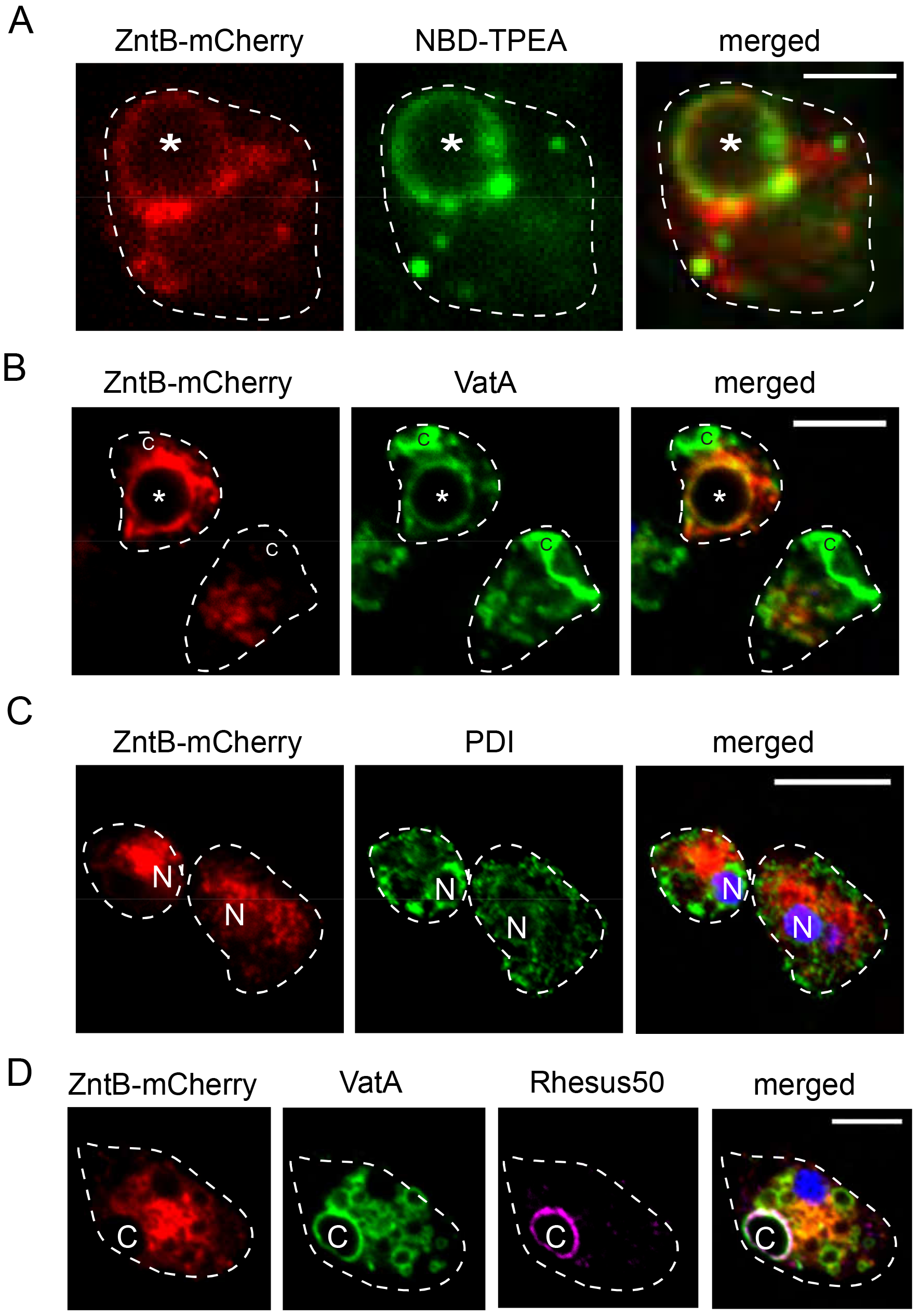
ZntB-mCherry decorates BCPs and does not co-localize with ER- and CV-markers. **A.** ZntB is present at BCPs at the lysosomal maturation stage. Cells expressing ZntB-mCherry were stained with NBD-TPEA and fed with 3 μm latex beads. Images were taken live. Scale bar, 5 μm. **B.** ZntB-mCherry co-localizes with the vATPase at BCPs. Scale bar, 10 μm. **C. and D.** ZntB-mCherry does not localize at the ER and the at the CV. ZntB-mCherry expressing cells were fixed and stained with antibodies against PDI (C) and Rhesus50 (D). Nuclei were stained with DAPI. Scale bars, 10 μm (C) and 5 μm (D). Asterisks label a BCPs, C: tubular network of the CV, N: nucleus.

**Fig. S4.**
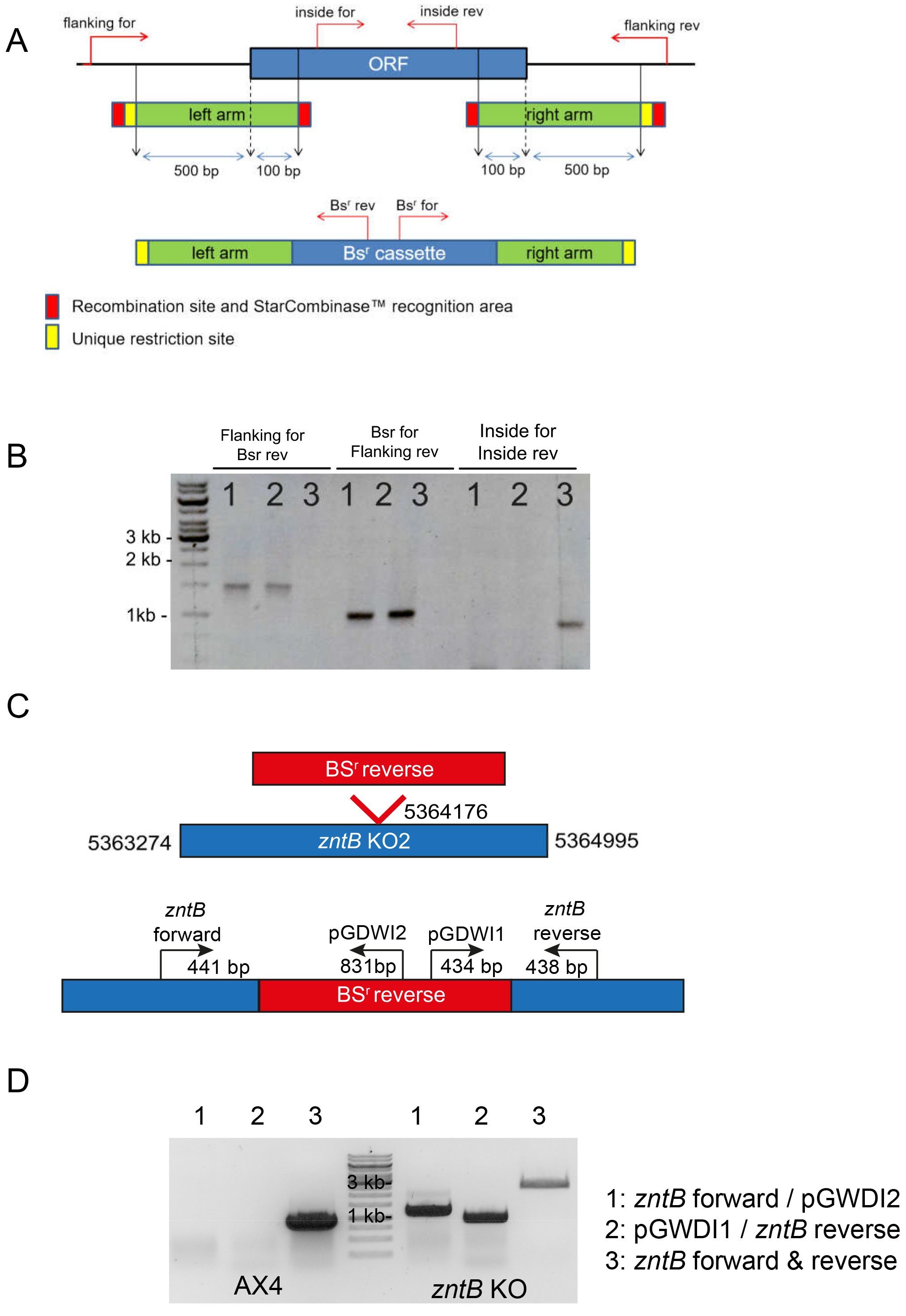
Generation of a *zntA* KO by homologous recombination and localization of the insertion in the *zntB* KO. **A.** Schematic drawing of the *zntA*-encoding gene locus (ORF, blue) flanked by non-coding segments. For gene disruption, the resistance cassette (BSr, green) was integrated removing a segment in themiddle of the gene (between the inside forward/inside reverse primers) using the StarCombinase and the StarGate cloning kit. The red arrows indicate primers that were used to monitor correct integration. **B.** PCR-analysis of two *zntA* mutants (#1 and #2) and wild type (#3). Using the flanking forward/BSR reverse or the flanking reverse/BSR forward primer combinations small products were obtained in both mutants, but not in the wild type. The inside forward/inside reverse primer combination yielded a small product in the wild type, but not in the mutants. Experiments were performed using mutant #1. **C.** The restriction-mediated insertion of the *zntB* KO interrupts the gene approximately in the middle at chromosomal position 5364176 (Chromosome 3). **D.** A diagnostic PCR was performed to confirm the insertion into *zntB* using the primers indicated in C.

**Fig. S5.**
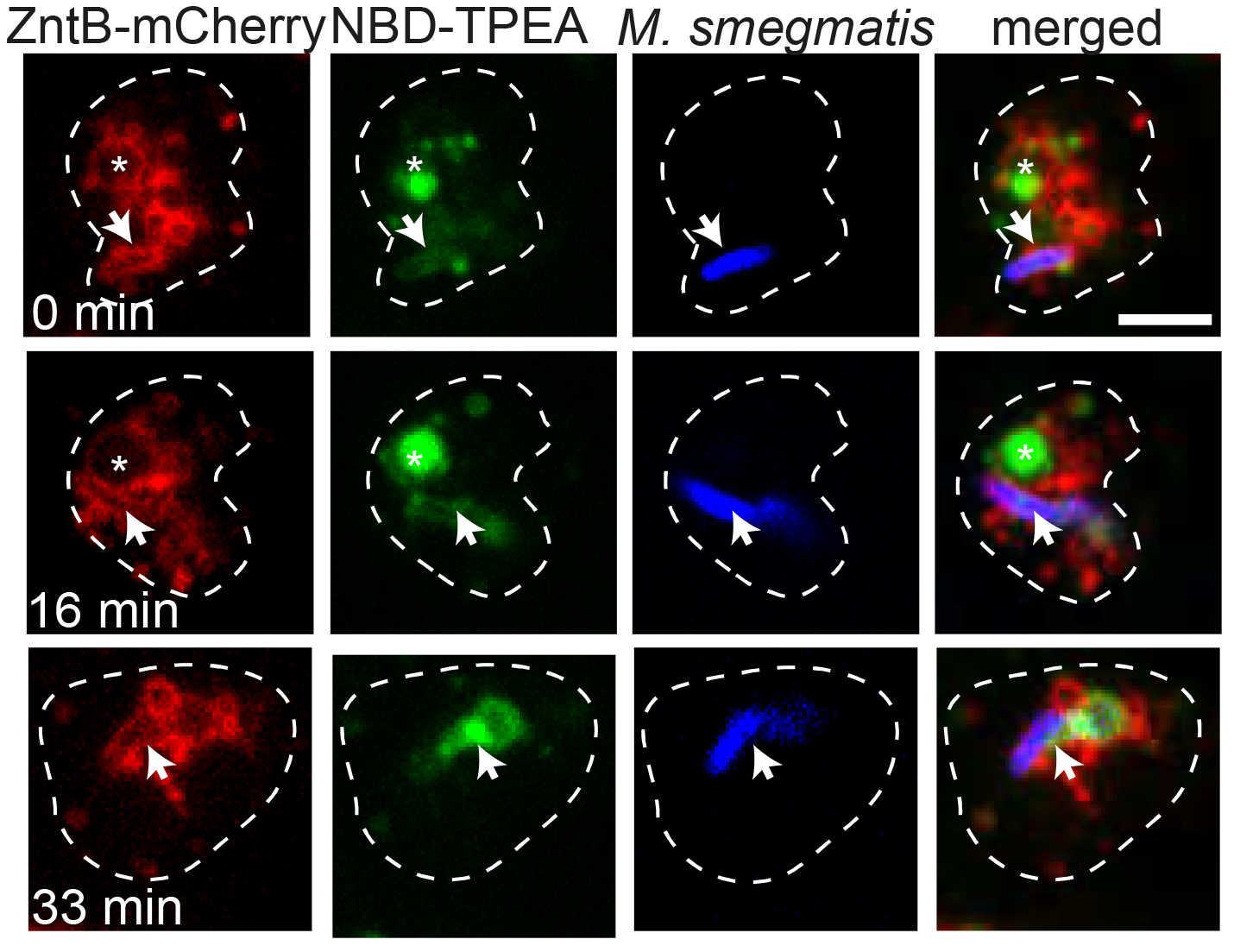
ZntB-mCherry is localized at the *M. smegmatis-containing* phagosome. Cells expressing ZntB-mCherry were stained with NBD-TPEA and fed with *M. smegmatis*. Images were recorded live. Scale bar, 5 μm. Arrows label phagosomes, asterisks point to zincosomes.

**Fig. S6.**
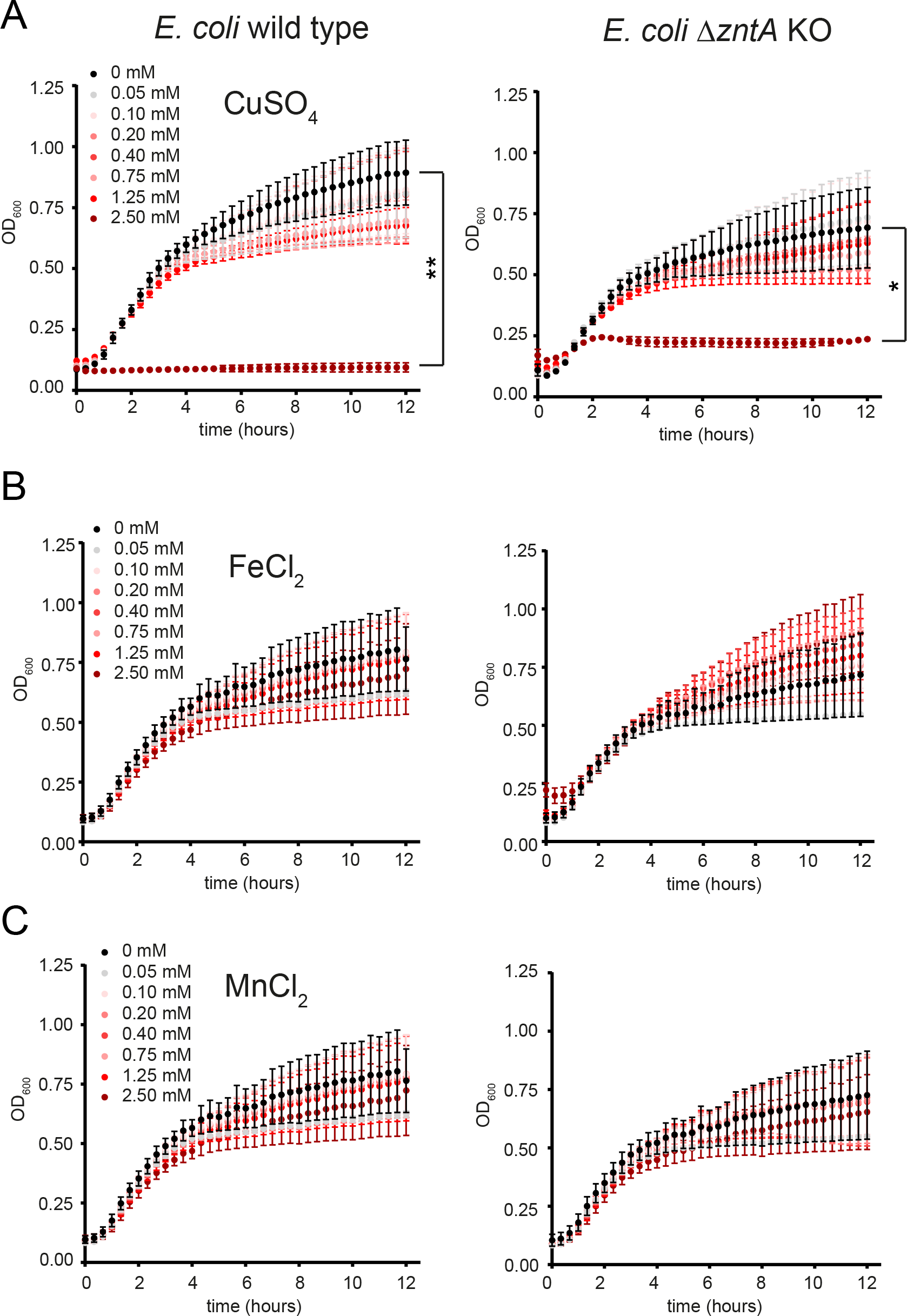
The *zntA E. coli* KO is not susceptible to increasing concentrations of CuSO_4_, FeCl_2_, MnCl_2_. *E. coli* strains were incubated in LB. Metals were added as indicated. The OD_600_ was measured with the help of a 96-well plate reader (SpectraMax i3, Molecular Devices). Statistical differences were calculated with a Bonferroni post hoc test after two-way ANOVA. Significantly different values were indicated by an asterisk (* P < 0.5, ** P < 0.01).

**Movie 1. Zn is expelled from the cells when the CV discharges.** For more information, see Fig. 1C.

**Movie 2. Dynamics of Zn in BCPs of NBD-TPEA-labelled cells.** For more information, see Fig. 2A.

**Movie 3. Dynamics of Zn in BCPs of FZ-3-labelled cells.** For more information, see Fig. 2C.

**Movie 4. Zincosome-BCP-fusion.** For more information, see Fig. 3A.

**Movie 5. Zincosome-BCP-fission.** For more information, see Fig. 3C.

**Movie 6. Dynamics of ZntB-mCherry at the BCP.** For more information, see Fig. 4D.

## References

Barisch, C., Lopez-Jimenez, A. T. and Soldati, T. (2015). Live Imaging of Mycobacterium marinum Infection in Dictyostelium discoideum. In Mycobacteria Protocols (Methods Mol Biol), vol. 1285 (eds T. Parish and D. Roberts), pp. 369–85: Humana, NYC.

Beard, S. J., Hughes, M. N. and Poole, R. K. (1995). Inhibition of the cytochrome bd-terminated NADH oxidase system in Escherichia coli K-12 by divalent metal cations. FEMS Microbiol Lett 131, 205–10.

Benghezal, M., Gotthardt, D., Cornillon, S. and Cosson, P. (2001). Localization of the Rh50-like protein to the contractile vacuole in Dictyostelium. Immunogenetics 52, 284–8.

Bird, A. J. (2015). Cellular sensing and transport of metal ions: implications in micronutrient homeostasis. J Nutr Biochem 26, 1103–15.

Bosomworth, H. J., Thornton, J. K., Coneyworth, L. J., Ford, D. and Valentine, R. A. (2012). Efflux function, tissue-specific expression and intracellular trafficking of the Zn transporter ZnT10 indicate roles in adult Zn homeostasis. Metallomics 4, 771–9.

Botella, H., Peyron, P., Levillain, F., Poincloux, R., Poquet, Y., Brandli, I., Wang, C., Tailleux, L., Tilleul, S., Charriere, G. M. et al. (2011). Mycobacterial p(1)-type ATPases mediate resistance to zinc poisoning in human macrophages. Cell Host Microbe 10, 248–59.

Bozzaro, S., Bucci, C. and Steinert, M. (2008). Phagocytosis and host-pathogen interactions in Dictyostelium with a look at macrophages. Int Rev Cell Mol Biol 271, 253–300.

Bozzaro, S., Buracco, S. and Peracino, B. (2013). Iron metabolism and resistance to infection by invasive bacteria in the social amoeba Dictyostelium discoideum. Front Cell Infect Microbiol 3, 50.

Bracco, E., Peracino, B., Noegel, A. A. and Bozzaro, S. (1997). Cloning and transcriptional regulation of the gene encoding the vacuolar/H+ ATPase B subunit of Dictyostelium discoideum. FEBS Lett 419, 37–40.

Buckley, C. M., Heath, V. L., Gueho, A., Dove, S. K., Michell, R. H., Meier, R., Hilbi, H., Soldati, T., Insall, R. H. and King, J. (2018). PIKfyve/Fab1 is required for efficient V-ATPase delivery to phagosomes, phagosomal killing, and restriction of Legionella infection. bioRxiv.

Buracco, S., Peracino, B., Andreini, C., Bracco, E. and Bozzaro, S. (2017). Differential Effects of Iron, Zinc, and Copper on Dictyostelium discoideum Cell Growth and Resistance to Legionella pneumophila. Front Cell Infect Microbiol 7, 536.

Burlando, B., Evangelisti, V., Dondero, F., Pons, G., Camakaris, J. and Viarengo, A. (2002). Occurrence of Cu-ATPase in Dictyostelium: possible role in resistance to copper. Biochem Biophys Res Commun 291, 476–83.

Carnell, M., Zech, T., Calaminus, S. D., Ura, S., Hagedorn, M., Johnston, S. A., May, R. C., Soldati, T., Machesky, L. M. and Insall, R. H. (2011). Actin polymerization driven by WASH causes V-ATPase retrieval and vesicle neutralization before exocytosis. J Cell Biol 193, 831–9.

Chan, H., Babayan, V., Blyumin, E., Gandhi, C., Hak, K., Harake, D., Kumar, K., Lee, P., Li, T. T., Liu, H. Y. et al. (2010). The p-type ATPase superfamily. J Mol Microbiol Biotechnol 19, 5–104.

Charette, S. J., Mercanti, V., Letourneur, F., Bennett, N. and Cosson, P. (2006). A role for adaptor protein-3 complex in the organization of the endocytic pathway in Dictyostelium. Traffic 7, 1528–38.

Colvin, R. A., Holmes, W. R., Fontaine, C. P. and Maret, W. (2010). Cytosolic zinc buffering and muffling: their role in intracellular zinc homeostasis. Metallomics 2, 306–17.

Djoko, K. Y., Ong, C. L., Walker, M. J. and McEwan, A. G. (2015). The Role of Copper and Zinc Toxicity in Innate Immune Defense against Bacterial Pathogens. J Biol Chem 290, 18954–61.

Ducret, V., Gonzalez, M. R., Scrignari, T. and Perron, K. (2016). OprD Repression upon Metal Treatment Requires the RNA Chaperone Hfq in Pseudomonas aeruginosa. Genes (Basel) 7.

Dunn, J. D., Bosmani, C., Barisch, C., Raykov, L., Lefrancois, L. H., Cardenal-Munoz, E., Lopez-Jimenez, A. T. and Soldati, T. (2017). Eat Prey, Live: Dictyostelium discoideum As a Model for Cell-Autonomous Defenses. Front Immunol 8, 1906.

Flannagan, R. S., Heit, B. and Heinrichs, D. E. (2015). Antimicrobial Mechanisms of Macrophages and the Immune Evasion Strategies of Staphylococcus aureus. Pathogens 4, 826–68.

Gee, K. R., Zhou, Z. L., Qian, W. J. and Kennedy, R. (2002). Detection and imaging of zinc secretion from pancreatic beta-cells using a new fluorescent zinc indicator. J Am Chem Soc 124, 776–8.

Gonzalez, M. R., Ducret, V., Leoni, S. and Perron, K. (2018). Pseudomonas aeruginosa zinc homeostasis: Key issues for an opportunistic pathogen. Biochim Biophys Acta.

Gotthardt, D., Warnatz, H. J., Henschel, O., Bruckert, F., Schleicher, M. and Soldati, T. (2002). High-resolution dissection of phagosome maturation reveals distinct membrane trafficking phases. Mol Biol Cell 13, 3508–20.

Hacker, U., Albrecht, R. and Maniak, M. (1997). Fluid-phase uptake by macropinocytosis in Dictyostelium. J Cell Sci 110 (Pt 2), 105–12.

Hagedorn, M., Neuhaus, E. M. and Soldati, T. (2006). Optimized fixation and immunofluorescence staining methods for Dictyostelium cells. Methods Mol Biol 346, 327–38.

Hagedorn, M. and Soldati, T. (2007). Flotillin and RacH modulate the intracellular immunity of Dictyostelium to Mycobacterium marinum infection. Cell Microbiol 9, 2716–33.

Hao, X., Luthje, F., Ronn, R., German, N. A., Li, X., Huang, F., Kisaka, J., Huffman, D., Alwathnani, H. A., Zhu, Y. G. et al. (2016). A role for copper in protozoan grazing - two billion years selecting for bacterial copper resistance. Mol Microbiol 102, 628–641.

Heuser, J., Zhu, Q. and Clarke, M. (1993). Proton pumps populate the contractile vacuoles of Dictyostelium amoebae. J Cell Biol 121, 1311–27.

Huang, L., Kirschke, C. P. and Gitschier, J. (2002). Functional characterization of a novel mammalian zinc transporter, ZnT6. J Biol Chem 277, 26389–95.

Kambe, T., Tsuji, T., Hashimoto, A. and Itsumura, N. (2015). The Physiological, Biochemical, and Molecular Roles of Zinc Transporters in Zinc Homeostasis and Metabolism. Physiol Rev 95, 749–84.

Kapetanovic, R., Bokil, N. J., Achard, M. E., Ong, C. Y., Peters, K. M., Stocks, C. J., Phan, M. D., Monteleone, M., Schroder, K., Irvine, K. M. et al. (2016). Salmonella employs multiple mechanisms to subvert the TLR-inducible zinc-mediated antimicrobial response of human macrophages. FASEB J.

Kehl-Fie, T. E. and Skaar, E. P. (2010). Nutritional immunity beyond iron: a role for manganese and zinc. Curr Opin Chem Biol 14, 218–24.

Kirschke, C. P. and Huang, L. (2003). ZnT7, a novel mammalian zinc transporter, accumulates zinc in the Golgi apparatus. J Biol Chem 278, 4096–102.

Kirsten, J. H., Xiong, Y., Davis, C. T. and Singleton, C. K. (2008). Subcellular localization of ammonium transporters in Dictyostelium discoideum. BMC Cell Biol 9, 71.

Kolaj-Robin, O., Russell, D., Hayes, K. A., Pembroke, J. T. and Soulimane, T. (2015). Cation Diffusion Facilitator family: Structure and function. FEBS Lett 589, 1283–95.

Krezel, A. and Maret, W. (2006). Zinc-buffering capacity of a eukaryotic cell at physiological pZn. J Biol Inorg Chem 11, 1049–62.

Leiba, J., Sabra, A., Bodinier, R., Marchetti, A., Lima, W. C., Melotti, A., Perrin, J., Burdet, F., Pagni, M., Soldati, T. et al. (2017). Vps13F links bacterial recognition and intracellular killing in Dictyostelium. Cell Microbiol.

Lopez, C. A. and Skaar, E. P. (2018). The Impact of Dietary Transition Metals on Host-Bacterial Interactions. Cell Host Microbe 23, 737–748.

Marchetti, A., Lelong, E. and Cosson, P. (2009). A measure of endosomal pH by flow cytometry in Dictyostelium. BMC Res Notes 2, 7.

McDevitt, C. A., Ogunniyi, A. D., Valkov, E., Lawrence, M. C., Kobe, B., McEwan, A. G. and Paton, J. C. (2011). A molecular mechanism for bacterial susceptibility to zinc. PLoS Pathog 7, e1002357.

Neuhaus, E. M., Almers, W. and Soldati, T. (2002). Morphology and dynamics of the endocytic pathway in Dictyostelium discoideum. Mol Biol Cell 13, 1390–407.

Neuhaus, E. M., Horstmann, H., Almers, W., Maniak, M. and Soldati, T. (1998). Ethane-freezing/methanol-fixation of cell monolayers: a procedure for improved preservation of structure and antigenicity for light and electron microscopies. J Struct Biol 121, 326–42.

Nies, D. H. (1999). Microbial heavy-metal resistance. Appl Microbiol Biotechnol 51, 730–50.

Ong, C. L., Gillen, C. M., Barnett, T. C., Walker, M. J. and McEwan, A. G. (2014). An antimicrobial role for zinc in innate immune defense against group A streptococcus. J Infect Dis 209, 1500–8.

Palmiter, R. D. and Findley, S. D. (1995). Cloning and functional characterization of a mammalian zinc transporter that confers resistance to zinc. EMBO J 14, 639–49.

Patrushev, N., Seidel-Rogol, B. and Salazar, G. (2012). Angiotensin II requires zinc and downregulation of the zinc transporters ZnT3 and ZnT10 to induce senescence of vascular smooth muscle cells. PLoS One 7, e33211.

Peracino, B., Buracco, S. and Bozzaro, S. (2013). The Nramp (Slc11) proteins regulate development, resistance to pathogenic bacteria and iron homeostasis in Dictyostelium discoideum. J Cell Sci 126, 301–11.

Peracino, B., Wagner, C., Balest, A., Balbo, A., Pergolizzi, B., Noegel, A. A., Steinert, M. and Bozzaro, S. (2006). Function and mechanism of action of Dictyostelium Nramp1 (Slc11a1) in bacterial infection. Traffic 7, 22–38.

Pyle, C. J., Azad, A. K., Papp, A. C., Sadee, W., Knoell, D. L. and Schlesinger, L. S. (2017). Elemental Ingredients in the Macrophage Cocktail: Role of ZIP8 in Host Response to Mycobacterium tuberculosis. Int J Mol Sci 18.

Ravanel, K., de Chassey, B., Cornillon, S., Benghezal, M., Zulianello, L., Gebbie, L., Letourneur, F. and Cosson, P. (2001). Membrane sorting in the endocytic and phagocytic pathway of Dictyostelium discoideum. Eur J Cell Biol 80, 754–64.

Rensing, C., Mitra, B. and Rosen, B. P. (1997). The zntA gene of Escherichia coli encodes a Zn(II)-translocating P-type ATPase. Proc Natl Acad Sci U S A 94, 14326–31.

Sattler, N., Monroy, R. and Soldati, T. (2013). Quantitative analysis of phagocytosis and phagosome maturation. Methods Mol Biol 983, 383–402.

Soldati, T. and Neyrolles, O. (2012). Mycobacteria and the intraphagosomal environment: take it with a pinch of salt(s)! Traffic 13, 1042–52.

Subramanian Vignesh, K., Landero Figueroa, J. A., Porollo, A., Caruso, J. A. and Deepe, G. S., Jr. (2013). Granulocyte macrophage-colony stimulating factor induced Zn sequestration enhances macrophage superoxide and limits intracellular pathogen survival. Immunity 39, 697–710.

Sunaga, N., Monna, M., Shimada, N., Tsukamoto, M. and Kawata, T. (2008). Expression of zinc transporter family genes in Dictyostelium. Int J Dev Biol 52, 377–81.

Uchikawa, T., Yamamoto, A. and Inouye, K. (2011). Origin and function of the stalk-cell vacuole in Dictyostelium. Dev Biol 352, 48–57.

Valdivia, R. H. and Falkow, S. (1997). Probing bacterial gene expression within host cells. Trends Microbiol 5, 360–3.

Veltman, D. M., Akar, G., Bosgraaf, L. and Van Haastert, P. J. (2009). A new set of small, extrachromosomal expression vectors for Dictyostelium discoideum. Plasmid 61, 110–8.

Vinkenborg, J. L., Nicolson, T. J., Bellomo, E. A., Koay, M. S., Rutter, G. A. and Merkx, M. (2009). Genetically encoded FRET sensors to monitor intracellular Zn2+ homeostasis. Nat Methods 6, 737–40.

Wagner, D., Maser, J., Lai, B., Cai, Z., Barry, C. E., 3rd, Honer Zu Bentrup, K., Russell, D. G. and Bermudez, L. E. (2005). Elemental analysis of Mycobacterium avium-, Mycobacterium tuberculosis-, and Mycobacterium smegmatis-containing phagosomes indicates pathogen-induced microenvironments within the host cell’s endosomal system. J Immunol 174, 1491–500.

Weiss, G. and Carver, P. L. (2018). Role of divalent metals in infectious disease susceptibility and outcome. Clin Microbiol Infect 24, 16–23.

Wiegand, S., Kruse, J., Gronemann, S. and Hammann, C. (2011). Efficient generation of gene knockout plasmids for Dictyostelium discoideum using one-step cloning. Genomics 97, 321–5.

Wienke, D., Drengk, A., Schmauch, C., Jenne, N. and Maniak, M. (2006). Vacuolin, a flotillin/reggie-related protein from Dictyostelium oligomerizes for endosome association. Eur J Cell Biol 85, 991–1000.

Xu, Z., Gunn-Hee, K., Su, J. H., Min, J. J., Chongmok, L., Injae, S. and Yoon, J. (2009). An NBD-based colorimetric and fluorescent chemosensor for Zn^2+^ and its use for detection of intracellular zinc ions. Tetrahedron 65, 2307–2312.

Yates, R. M., Hermetter, A. and Russell, D. G. (2005). The kinetics of phagosome maturation as a function of phagosome/lysosome fusion and acquisition of hydrolytic activity. Traffic 6, 413–20.

